# Decomposing tissue-of-origin aging clocks reveals composite aging states across cancers

**DOI:** 10.64898/2026.11.12.765990

**Authors:** Payton Bock, Jackson Smith, Pranav Narnur, Nageen Zahra, Nuzulul Kurniansyah, Stefano Monti

**Affiliations:** Section of Computational Biomedicine, Boston University Chobanian & Avedisian School of Medicine, Boston, MA 02118, USA; Bioinformatics Program, Center for Computing & Data Sciences, Boston University, Boston, MA 02215, USA; Graduate Program in Genetics & Genomics, Boston University Chobanian & Avedisian School of Medicine, Boston, MA 02118, USA; Section of Biomedical Genetics, Boston University Chobanian & Avedisian School of Medicine, Boston, MA 02118, USA; Department of Biostatistics, Boston University School of Public Health, Boston, MA 02118, USA

## Abstract

Aging is accompanied by tissue-specific molecular remodeling, yet whether tumors retain or depart from these programs remains unclear. In a pan-cancer analysis we show that age-associated molecular changes in tumors are organized by tissue of origin and partially mirror matched normal tissue trends. We introduce tissue-of-origin aging clocks to measure each tumor’s deviation from its expected tissue aging, and *ClockSHAP* to decompose deviations into pathway-level contributions. Across breast and five other cancers, this decomposition reveals composite aging states in which youth-like deviations in proliferative pathways coexist with older-like deviations in damage-response and signaling pathways. In breast cancer, a clock trained only on normal tissues, applied without retraining, reproduces its deviations and pathway structure in two independent cohorts (∼9,700 tumors), aligns with molecular subtype and *TP53* mutation status, and predicts survival. These findings show how cancers reshape tissue-of-origin aging programs and provide a framework for applying and interpreting aging clocks in disease.

## Introduction

Chronological age is the dominant risk factor for most cancers^1^, and many biological hallmarks of aging (e.g., genomic instability, epigenetic alteration, chronic inflammation) overlap with, and are frequently co-opted by, tumorigenesis.^2^ Normal tissues undergo tissue-specific age-related changes in cancer-relevant programs, including immune, proliferative, metabolic, and stress-response pathways.^3,4^ Tumors therefore arise within tissues that are themselves aging.

Efforts to characterize the cancer–aging relationship have largely focused on tumors themselves. Pan-cancer studies have reported age-associated patterns spanning multiple molecular layers.^5–8^ These studies establish that cancers vary molecularly with age but generally characterize age-associated changes within cancers rather than relative to the aging of their tissue of origin. Without a normal aging reference, it remains unclear whether tumor molecular states preserve, reshape, or depart from those aging programs.

Aging clocks, models that estimate age from molecular profiles, offer an elegant approach to make such a comparison. The gap between a clock’s predicted age and chronological age is often summarized as age acceleration and used as a readout of a molecular aging state.^9^ Clocks have been applied to cancers to ask whether tumors carry altered molecular-age signals, typically by summarizing each sample as a predicted molecular age or age-acceleration value and relating that value to molecular and clinical features including mutation burden, stage, and survival.^8,10–14^ Such findings have motivated hypotheses that aging clocks may capture processes such as epigenetic maintenance^11^, epigenetic drift and senescence^8^, and stemness or cellular plasticity.^10^

However, what these values capture in cancer remains unclear. Clock outputs are often treated as black boxes: differences between predicted and chronological age are associated with tumor features and outcomes, but the feature-level biology underlying those differences is often not resolved.^9^ In cancer, this limitation is compounded because tumors can appear younger or older depending on the clock and feature space used, and a single predicted-age or age-acceleration value does not reveal what the clock is capturing about tumor aging biology.^8,10–15^

Here, we ask whether tumors depart from the aging expected of their tissue of origin, and in what biological directions. The study proceeds in two phases. In the first, we characterize age-associated molecular changes across 33 cancer types and six data modalities – somatic mutations, copy-number alterations, DNA methylation, transcriptomics, microRNAs, and RPPA-based proteomics – from The Cancer Genome Atlas (TCGA)^16^, to establish that tumor aging variation is structured by tissue of origin, motivating a tissue-specific reference frame for subsequent analyses.

In the second, we focus on transcriptomics, which affords the harmonized multi-cohort data needed to train interpretable, pathway-level aging clocks on normal tissue and to apply them without retraining across independent tumor cohorts. We compare transcriptomic aging trends between cancers and corresponding normal tissues in the Genotype-Tissue Expression (GTEx) project^17^, finding tissue-dependent partial preservation of normal aging programs. Because shared aging trends do not necessarily mean a shared molecular state, we next train tissue-of-origin aging clocks in GTEx and quantify each tumor’s deviation from its age-matched normal expectation. We introduce ClockSHAP, an interpretability method for aging clocks that decomposes each deviation into feature-level contributions. We apply the framework to six cancers, with particular focus on breast cancer, where large independent cohorts are available for replication. This reveals composite aging states in which youth-like proliferative contributions often coexist with older-like damage-response and signaling contributions. In breast cancer, these states align with established tumor biology and replicate in two large independent cohorts. Together, these results show how cancers reshape the aging programs of their tissue of origin and provide a framework for determining what aging clocks capture when applied to disease.

## Results

### Age-associated molecular changes in tumors are heterogeneous but structured by tissue of origin

We first asked how extensively tumor molecular profiles vary with patient age. Across 33 TCGA cancer types, we identified age-associated features in six molecular layers: DNA methylation, RNA-seq, miRNA, RPPA, somatic mutation, and somatic copy-number alteration (SCNA). We analyzed each cancer–layer combination containing at least 50 samples with complete covariate data. The number of significant age-associated features varied substantially across cancer types and molecular layers (FDR ≤ 0.05; Fig. 1a), although absolute counts are not directly comparable across layers because they differ in feature number and resolution. The complete per-feature results underlying Fig. 1a can be browsed in the Pan-Cancer Aging Explorer (https://montilab.github.io/pancancer-aging-explorer/). Within each layer, both the breadth and direction of age-associated change varied markedly among cancers. Tumors therefore showed widespread but heterogeneous molecular differences with patient age rather than a uniform pan-cancer aging pattern.

**Figure 1.**
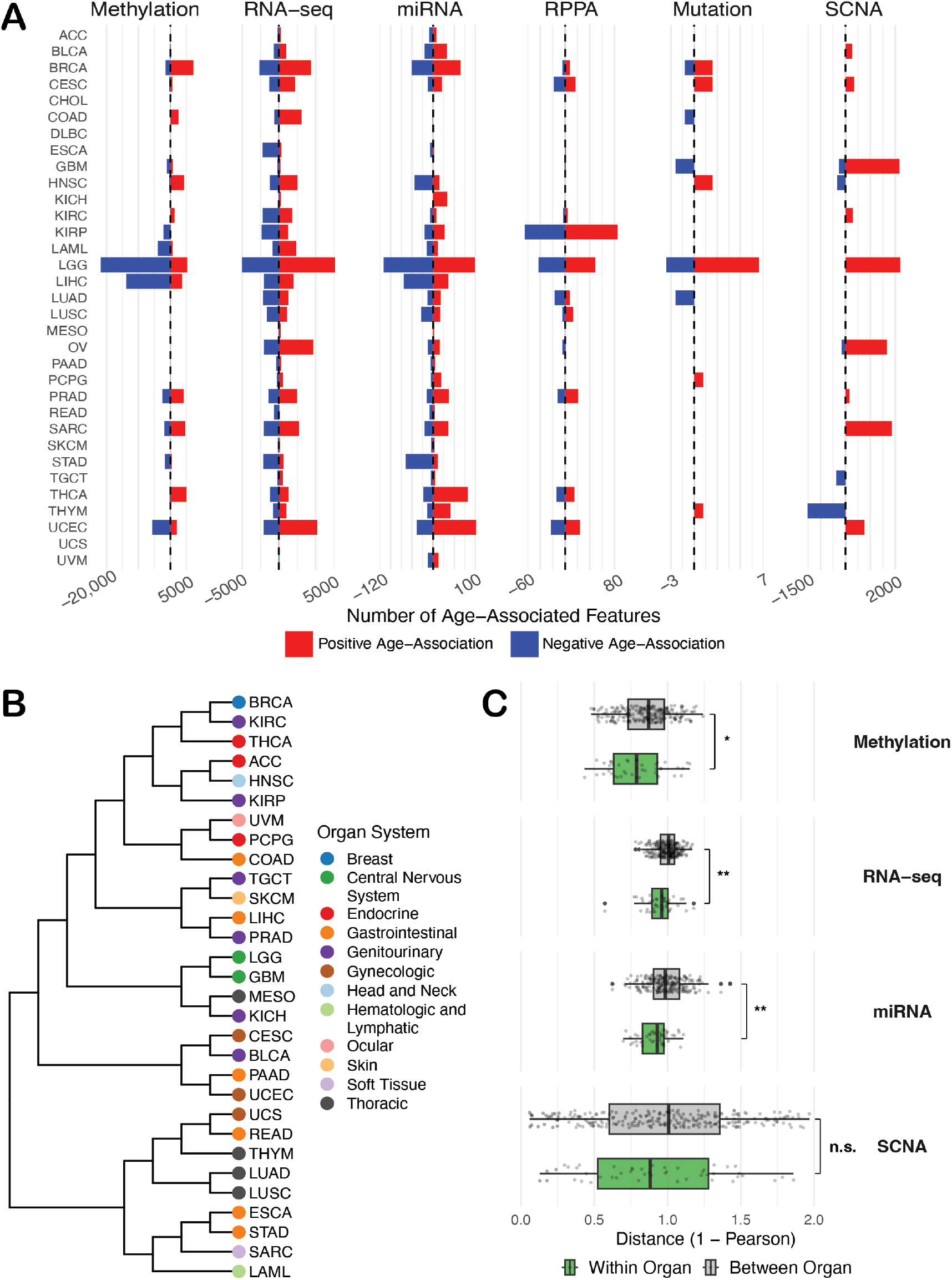
Pan-cancer molecular age associations and their tissue-of-origin structure across TCGA. (a) Number of significant age-associated features (Benjamini-Hochberg adjusted P ≤ 0.05) per TCGA cancer type (n = 33) across six molecular layers: DNA methylation (differentially methylated regions), RNA-seq, miRNA, RPPA, somatic mutation, and somatic copy-number alteration (SCNA). Bars are colored by direction of association (red, increased with age; blue, decreased with age). Counts are at the region level for methylation, gene level for somatic mutation and SCNA, and feature level otherwise. Note the differing count-axis scales across layers. (b) Hierarchical clustering of TCGA cancers (n = 30) on the cross-cancer similarity of per-feature age-effect profiles, averaged across methylation, RNA-seq, miRNA, and SCNA (1 − Pearson); RPPA and somatic mutation were excluded, and clustering was restricted to cancers with results in both methylation and RNA-seq (see Methods). Tip points are colored by tissue-of-origin organ system. (c) Pairwise distances (1 − Pearson) between cancer types, split into within-organ-system (green) and between-organ-system (grey) pairs and shown per layer. Each layer is evaluated on the cancers assayed in that layer, so the number of pairs and the number of retained organ systems differ between panels; single-cancer organ systems were excluded within each layer. Boxes show the median and interquartile range, whiskers extend to 1.5× the interquartile range, and points are individual cancer-type pairs. Within-organ versus between-organ distances were compared per layer by permutation of organ-system labels across cancers (10,000 permutations, two-sided): methylation P = 0.034 (n = 43 within, 233 between), RNA-seq P = 6 × 10⁻⁴ (46, 254), miRNA P = 0.0026 (45, 208), and SCNA P = 0.28 (48, 252).

We next asked whether this heterogeneity was organized by tissue of origin. For each molecular layer, we represented each cancer type by its profile of per-feature age effects and calculated the similarity of these profiles across cancer types. Four layers had sufficient cross-cancer feature overlap for this analysis: DNA methylation, RNA-seq, miRNA, and SCNA. RPPA and somatic mutation were excluded because too few features were shared across cancers (Methods). Integrating age-effect profiles across these layers revealed tissue-of-origin structure among 30 cancer types (CHOL, DLBC, and OV lacked results in methylation or RNA-seq and were not clustered). Hierarchical clustering showed that cancers arising within the same organ system frequently grouped together. Prominent examples included tight pairings of the central nervous system cancers GBM and LGG and the gastrointestinal cancers ESCA and STAD. Three thoracic cancers also formed a local cluster, with LUAD and LUSC paired most closely (Fig. 1b). These patterns indicated that related tissues share aspects of how their tumor molecular profiles vary with age, similar to the known organization of tumor molecular profiles by cell of origin.^18^

To quantify this pattern, we compared pairwise distances between cancers from the same organ system with distances between cancers from different organ systems. Within-organ-system cancers had more similar age-effect profiles for DNA methylation (P = 0.034), RNA-seq (P = 6 × 10⁻⁴), and miRNA (P = 0.0026). Within-organ distances were also lower for SCNA, though not significantly so (P = 0.28; Fig. 1c). Age-associated tumor changes were therefore heterogeneous but not unstructured, carrying tissue-of-origin organization across several molecular layers and establishing tissue of origin as a relevant context for interpreting tumor aging.

### Tumors partially preserve normal tissue aging programs in a tissue-specific manner

Because age-associated tumor changes were structured by tissue of origin, we next asked to what extent they reflected aging programs observed in normal tissues. We compared age-associated transcriptomic changes between GTEx normal tissues and TCGA tumors using two complementary readouts: Hallmark pathways and estimated cell-type composition. For the pan-tissue analysis, we estimated age associations within each GTEx tissue and TCGA cancer, restricting the analysis to cohorts with at least 50 samples. This yielded 27 normal tissues and 31 tumor types, whose effects were meta-analyzed separately (Fig. 2a,b).

**Figure 2.**
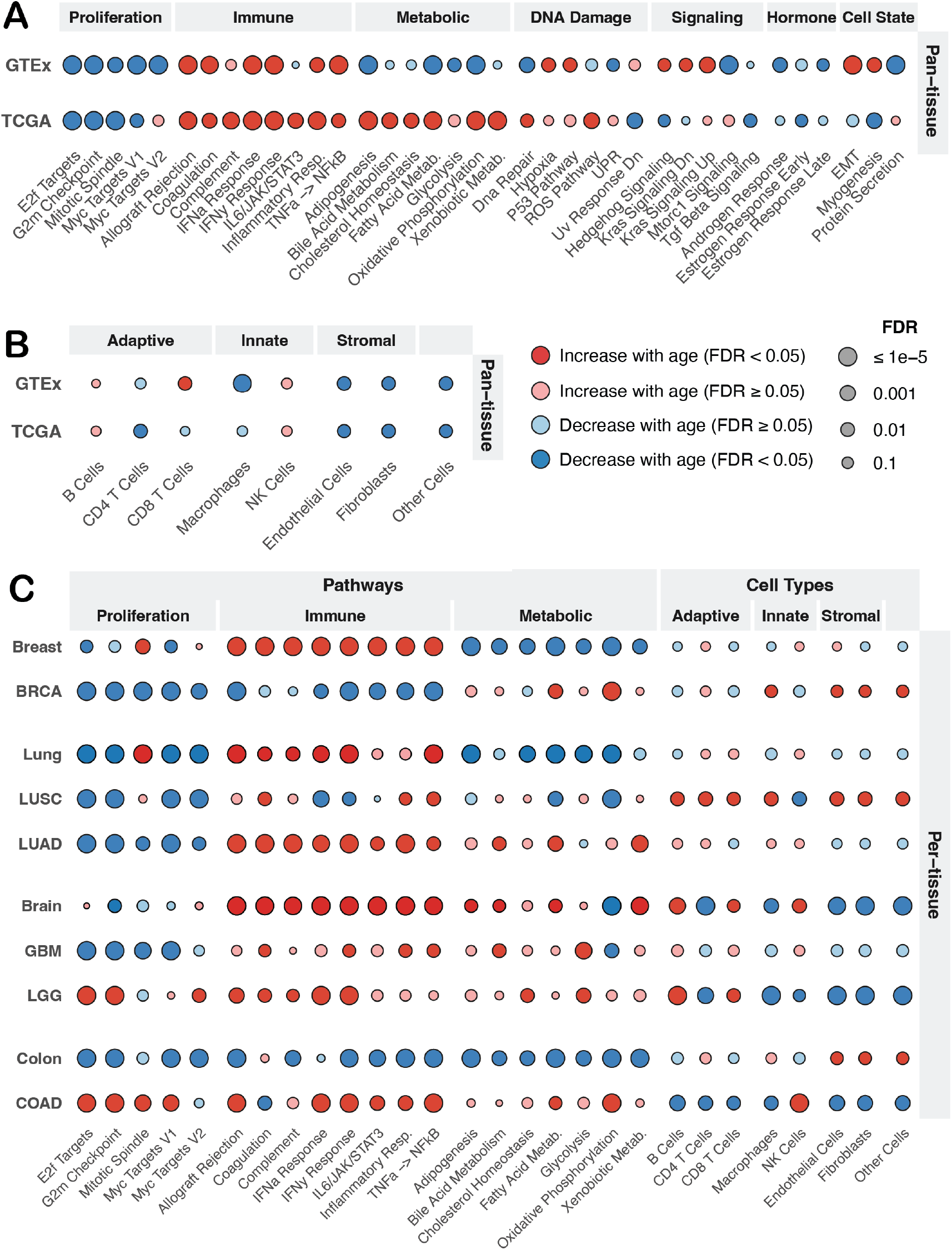
Cross-cohort agreement of age-associated pathway and cell-type programs between GTEx and TCGA. (a) Pan-tissue agreement in Hallmark pathway aging between normal tissue (GTEx, n = 27 tissues) and tumors (TCGA, n = 31 cancers), from within-cohort random-effects meta-analysis of per-project gene-level age associations. Pathways with FDR < 0.05 in at least one cohort are shown and grouped into broad functional categories for visualization. (b) Pan-tissue agreement in cell-type compositional aging between GTEx and TCGA (same cohorts as in a), from within-cohort meta-analysis of per-project compositional age effects, with cell types grouped into adaptive, innate, stromal, and other compartments. (c) Tissue-matched agreement for the four emphasis tissues (breast, BRCA; lung, LUAD and LUSC; brain, GBM and LGG; colon, COAD), showing pathway programs (restricted to proliferation, immune, and metabolic categories) and cell-type programs side by side, with GTEx as the tissue-of-origin reference and TCGA the matched tumor cohorts. Across all panels, dot color indicates the direction and significance of the age association (red, increase with age; blue, decrease with age; darker, FDR < 0.05; lighter, FDR ≥ 0.05) and dot size indicates FDR, with larger dots denoting stronger significance (legend in b). Cell-type proportions were estimated with EPIC, whose reference is derived from blood and tumor-microenvironment data rather than normal solid tissue; fibroblast estimates use the cancer-associated-fibroblast reference.

When organized into broad functional categories, normal tissues and tumors agreed on some aging programs and diverged on others at the pan-tissue level (Fig. 2a). Proliferative programs decreased with age in both cohorts, while immune programs, including interferon-related programs and inflammatory response, increased with age in both. Metabolic programs diverged consistently, decreasing with age in normal tissues but increasing with age in tumors, with adipogenesis, fatty acid metabolism, and oxidative phosphorylation significant in both cohorts. Pathway-level aging was therefore neither uniformly preserved nor reversed in cancer.

Age-associated changes in estimated cell-type composition were also largely directionally concordant between normal tissues and tumors, including decreases in fibroblasts and endothelial cells, whereas CD8 T cells showed the clearest directional divergence, increasing with age in normal tissues while decreasing in tumors (Fig. 2b). Together, the pathway and compositional analyses showed that tumors retained several features of normal tissue aging while departing from others.

Given the tissue-of-origin organization observed in tumor aging profiles, we next asked whether agreement with normal aging was itself tissue specific. We focused on four tissues with clear correspondence between a GTEx tissue and a TCGA cancer and sufficient samples in both cohorts, which define the emphasis set carried through the remainder of the study. We compared GTEx breast, lung, brain, and colon with the corresponding TCGA cancers BRCA, LUAD and LUSC, GBM and LGG, and COAD, respectively, focusing on the three program categories with the clearest pan-tissue patterns (Fig. 2c). Concordance at the per-tissue level was partial and varied across tissues. The age-related decline in proliferative programs was preserved across most tissue–tumor comparisons, with LGG and COAD standing out as exceptions in which several proliferative programs instead increased with age. Both are cancers whose molecular subtypes are themselves strongly age-associated, including IDH-defined glioma subtypes and early- versus late-onset colorectal cancer.^8,19^

Agreement with normal aging was not fixed by tissue of origin alone. LUAD and LUSC, both referenced to normal lung, differed in how closely they tracked it: LUAD broadly preserved the proliferative and immune trends of normal lung while diverging in metabolic programs, whereas LUSC showed opposing interferon trends and more substantial divergence in estimated cell-type composition. Among the four tissue groups, brain tumors showed comparatively strong agreement, whereas COAD and BRCA showed broad divergence from their corresponding normal tissues (Fig. 2c). Tumors therefore mirrored aspects of tissue-of-origin aging while reshaping or reversing others.

### Tissue-of-origin aging clocks quantify aging states and ClockSHAP decomposes them

The preceding analyses compared age-associated trends between cohorts, but a shared trend does not necessarily imply a shared molecular state. A tumor can change with age in the same direction as normal tissue while remaining shifted from the state expected in normal tissue at the same age. We use the term *aging state* to refer to the molecular configuration – summarized here by the profile of pathway activity scores – that characterizes a tissue at a given chronological age. An aging clock trained on normal tissue maps this configuration to a predicted age, and the gap between a tumor’s predicted age and the predicted age expected for normal tissue at the same chronological age quantifies how far the tumor’s aging state has shifted. We therefore next asked where individual tumors were positioned relative to this expectation. Because agreement with normal aging varied across tissues, this position had to be measured against the corresponding tissue of origin rather than a pooled normal reference.

We trained transcriptomic aging clocks in GTEx breast, brain, lung, and colon, each predicting chronological age from 50 Hallmark pathway scores (ssGSEA), establishing a normal aging reference for each tissue before applying the clocks to the corresponding tumors (Methods). Training R² was 0.17–0.26 across these four tissues; nested cross-validation yielded out-of-sample R² of 0.13–0.17 with RMSEs of 9.7–12.3 years, indicating that the 50-pathway clocks captured age-related signal in held-out samples (Supplementary Table 1; Supplementary Fig. 1).

We quantify each sample’s position relative to the normal aging expectation as the age-matched deviation: the difference between its clock-predicted age and the predicted age expected for a normal tissue sample of the same chronological age (Methods). Positive deviations indicate samples that appear older than expected for their age; negative values indicate samples that appear younger. This measure generalizes *relative age acceleration* (RAA)^9,20^ by carrying the normal-tissue reference forward when the clock is applied to new cohorts, so that tumors are measured against normal tissue rather than against one another.

To resolve what the clock captures beneath this summary value, we developed ClockSHAP, an interpretability method for linear-additive aging clocks that decomposes each deviation into exact feature-level contributions (Methods). Because our clocks were linear-additive and trained on Hallmark pathway scores, these contributions were pathway level and summed exactly to each sample’s total age-matched deviation.

We illustrate the framework with the lung clock applied to its own GTEx training cohort, highlighting two samples with deviations of similar magnitude but opposite sign (Fig. 3a). ClockSHAP revealed that the same pathway – epithelial-mesenchymal transition (EMT) – contributed in opposite directions for these two samples because they fell on opposite sides of its age-expected trend (Fig. 3b), and that the full combination of pathway contributions differed between them (Fig. 3c,d), demonstrating that contributions are sample-specific in both magnitude and direction. We interpret positive pathway contributions as “older-like” – shifting the prediction toward an older age relative to the age-matched expectation – and negative contributions as “youth-like”.

**Figure 3.**
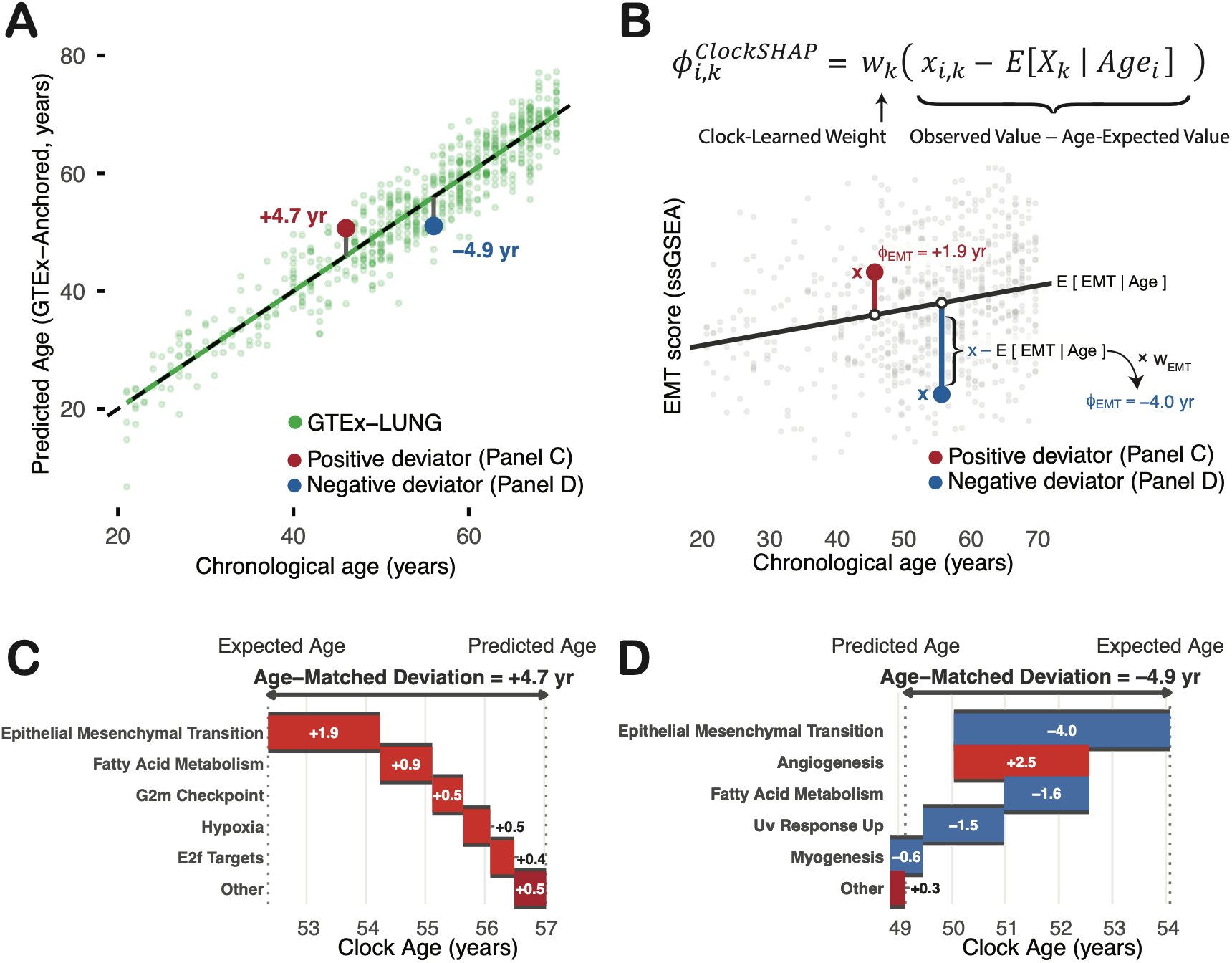
ClockSHAP decomposes age-matched deviation into pathway contributions, illustrated on GTEx lung. Demonstration of the framework using the lung clock, an elastic-net model predicting chronological age from 50 Hallmark ssGSEA pathway scores, applied to its own GTEx training cohort (n = 655 samples), where the mean deviation is zero, so the panels isolate how individual samples depart from the age-matched expectation. (a) GTEx-anchored predicted age (chronological age plus age-matched deviation) versus chronological age for GTEx lung samples; the reference is anchored to the identity line (dashed), so vertical distance from it equals the deviation. Two representative samples are highlighted, a positive deviator (red, +4.7 years) and a negative deviator (blue, −4.9 years). (b) The per-pathway mechanism, shown for epithelial-mesenchymal transition (EMT), with the ClockSHAP expression annotated above the panel. The line is the GTEx age-conditioned expectation E[EMT | Age], open circles mark that expectation and filled circles the observed score, and a pathway’s contribution is the clock weight times the sample’s departure from the line, φ = w (observed − age-expected). The two highlighted samples sit on opposite sides of the line, so the same pathway contributes positively for one and negatively for the other. (c) ClockSHAP waterfall for the positive deviator, decomposing its +4.7-year deviation into additive pathway contributions that carry the expected age to the predicted age; the five largest-magnitude contributors are shown individually and the remainder aggregated as “Other.” (d) ClockSHAP waterfall for the negative deviator, decomposing its −4.9-year deviation, displayed as in c. Contrasting c and d shows the decomposition is sample-specific in both magnitude and direction

### Tumor age-matched deviations vary across cancers and associate with survival

Having established the tissue-of-origin clock framework in GTEx, we next applied the four normal-tissue clocks to six corresponding TCGA cancers: BRCA, LUAD, LUSC, GBM, LGG, and COAD. For each tumor, the clock and age-conditioned feature expectations from its tissue of origin were retained as fixed references, allowing its predicted age to be compared with that expected for a normal sample of the same chronological age. Five of the six cancers showed substantial negative mean deviations, indicating youth-like positioning relative to their tissue-of-origin references (Fig. 4a). BRCA showed the largest mean deviation (−22.9 years), followed by LUSC (−18.8), GBM (−16.3), LUAD (−14.6), and COAD (−9.4), whereas LGG was centered near its normal-tissue expectation (−0.1 years). Therefore, the magnitude of departure from normal tissue aging varied markedly across cancers but was predominantly youth-like. Within each cancer, deviation became more youth-like as tumor purity increased, as expected if the youth-like shift arises in tumor cells and is diluted by surrounding non-tumor tissue (Supplementary Fig. 2a).

**Figure 4.**
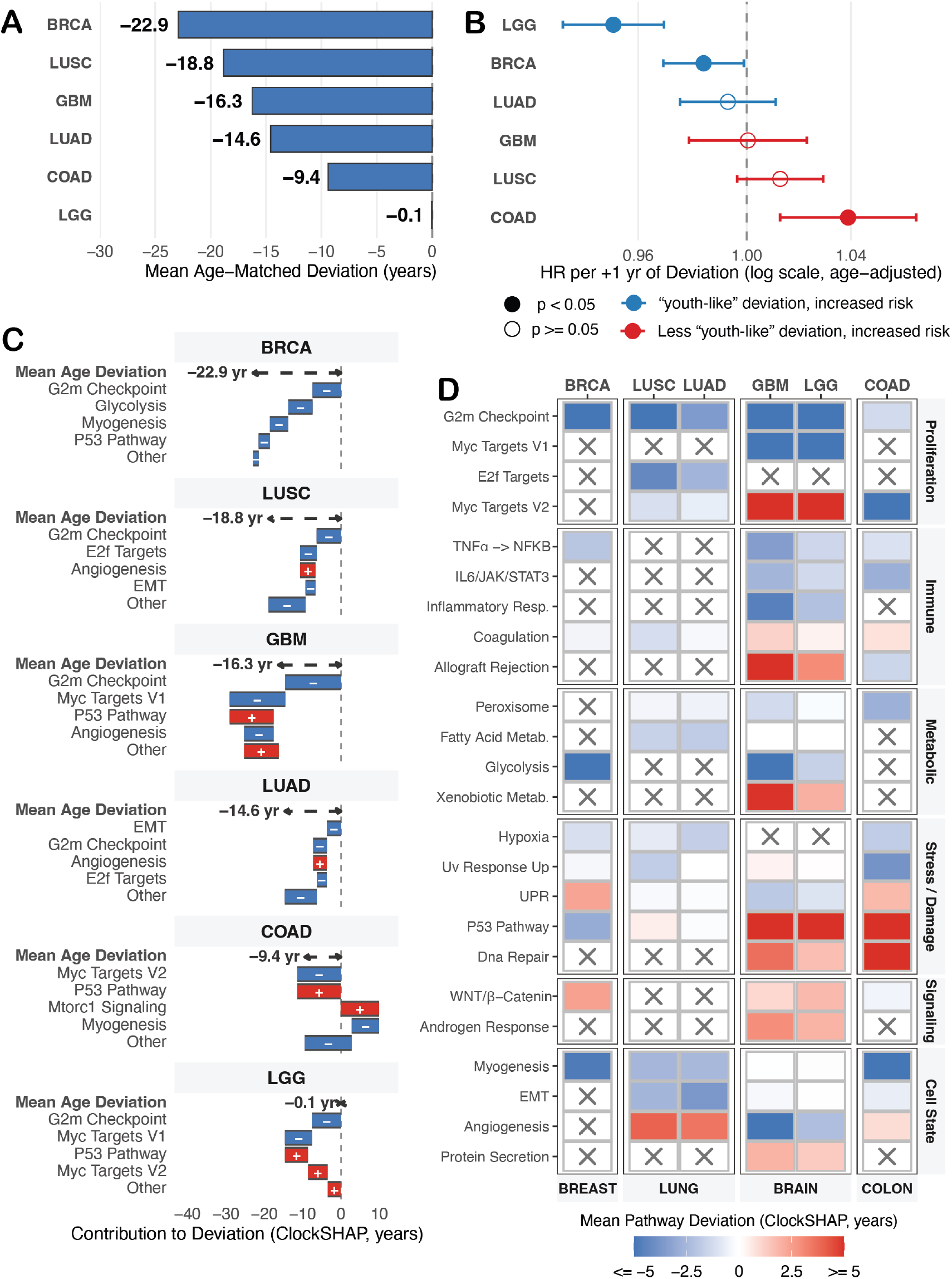
Tumors show large, tissue-structured age-matched deviations that decompose into recurring pathway programs. Cross-cancer analysis of the six emphasis cancers, each scored with its tissue-of-origin clock and ClockSHAP. (a) Mean age-matched deviation per cancer, ordered from most to least youth-like; five of six cancers deviate substantially youth-like from their tissue-of-origin reference (BRCA −22.9, LUSC −18.8, GBM −16.3, LUAD −14.6, COAD −9.4 years), with LGG near zero (−0.1). (b) Age-adjusted Cox hazard ratio for overall survival per additional year of deviation, one model per cancer (n = 156 to 1,116 tumors, 109 to 210 events per cancer); error bars are 95% confidence intervals on a log scale, point color indicates the direction of risk (blue, more youth-like deviation associates with higher risk; red, less youth-like with higher risk), and filled points mark p < 0.05 (open, p ≥ 0.05). Deviation was associated with survival in LGG (HR 0.95, p = 2 × 10⁻⁷), COAD (HR 1.04, p = 0.003), and BRCA (HR 0.98, p = 0.04). (c) Decomposition of each cancer’s mean deviation (annotated per panel, as in a) into additive pathway contributions (ClockSHAP), showing the top four pathways per cancer and the aggregate of the remainder (“Other”). (d) Mean per-pathway deviation across cancers, restricted to pathways whose mean contribution reaches at least one year in at least two cancers, grouped by pathway category (rows) and tissue of origin (columns); color is the mean contribution in years (capped at ±5), and a grey cross marks pathways not retained by that cancer’s clock.

Adjacent normal tissue, available for four of the six cancers, provided a within-TCGA comparator. In BRCA, LUSC, and LUAD, adjacent normal deviated less than the matched tumors (−5.3, −7.4, and −8.8 years against −22.9, −18.8, and −14.6), positioning it between the GTEx reference and the tumor and indicating that the tumor deviations did not arise from a uniform offset between cohorts. COAD was the exception, with adjacent normal deviating more than its tumors (−11.7 against −9.4; Supplementary Table 2). A prior transcriptomic comparison of GTEx normal, TCGA adjacent normal, and TCGA tumors across eight tissues reported this same structure^21^, with adjacent normal occupying an intermediate state between normal tissue and tumor. Colon was the sole exception, with adjacent normal instead resembling tumor, which that study attributed to anatomical heterogeneity in the GTEx colon dataset.

We also asked whether these deviations captured clinically relevant variation, as age-acceleration values are commonly related to outcomes in aging-clock studies.^9,12,13,22^ In age-adjusted Cox models, deviation was associated with overall survival in LGG, COAD, and BRCA, although the direction of association differed across cancers (Fig. 4b). In LGG, the proportional hazards assumption did not hold, and the reported estimate therefore represents a time-averaged effect (Supplementary Table 3). In LGG and BRCA, more youth-like deviations were associated with higher risk, whereas in COAD, less youth-like deviations were associated with higher risk. These associations were unchanged when tumor purity was added to the models (Supplementary Table 4). Together, these findings suggested that a tumor’s position relative to its normal aging reference captured clinically relevant variation in a subset of cancers, although the direction of that relationship was cancer specific.

### ClockSHAP reveals composite tumor aging states across cancers

To resolve which clock features drove these deviations, we decomposed each tumor’s age-matched deviation into pathway-level contributions with ClockSHAP. We then summarized each cancer by the mean contribution of each pathway, highlighting the four largest-magnitude contributors and aggregating the remainder as “Other” (Fig. 4c).

The pathways contributing most strongly to these deviations were recurrent and recognizable cancer programs (Fig. 4c). G2M checkpoint was among the top contributors in five of six cancers and was youth-like in each, while other proliferative programs, including Myc targets and E2F targets, frequently contributed in the same direction. Beyond this shared proliferative signal, the leading contributors that distinguished cancers ran in both directions. EMT was youth-like in the lung cancers, and glycolysis was youth-like in BRCA. In the other direction, P53 pathway was older-like in GBM, LGG, and COAD, angiogenesis was older-like in the lung cancers, and mTORC1 signaling was older-like in COAD. Thus, although the clocks were trained only on normal tissue, the mean deviation in each cancer resolved into contributions from canonical cancer-associated programs.

To identify broader directional themes, we focused on pathways with an absolute mean contribution of at least one year in two or more cancers (Fig. 4d). Proliferative programs were youth-like in nearly all cancers, with G2M checkpoint youth-like throughout. The main exception was Myc Targets V2, which was older-like in the brain cancers. Immune programs, where retained, also tended to be youth-like, including TNFα–NFκB signaling, inflammatory response, and IL6/JAK/STAT3 signaling. Among metabolic programs, glycolysis contributed in the youth-like direction in BRCA, GBM, and LGG. Older-like contributions were concentrated in damage-response and signaling programs: P53 pathway was older-like in every cancer except BRCA, DNA repair was older-like in the brain and colon cancers, Wnt/β-catenin was older-like in BRCA and the brain cancers, and androgen response was older-like in the brain cancers. Cell-state programs were mixed, with myogenesis and EMT youth-like where retained, and angiogenesis among the strongest older-like signals in the lung cancers.

These themes were shared across cancers, but the decomposition was also tissue- and cancer-specific. Each tissue-of-origin clock retained a different set of pathways, meaning that some programs were excluded from tumor types whose corresponding clock did not use them (grey crosses, Fig. 4d). Additional pathways that did not meet the cross-cancer recurrence threshold nevertheless made substantial contributions in individual cancers (Supplementary Fig. 3). This pattern echoed the tissue-specific organization of aging observed across cancers and normal tissues (Figs. 1 and 2).

Taken together, these decompositions showed that a tumor’s net deviation did not represent a uniform shift toward a younger or older state, but rather the balance of opposing pathway contributions. Youth-like proliferative programs pulled tumors below their normal-tissue expectation, while older-like damage-response and signaling programs often pulled in the opposite direction. This structure was clearest in LGG, whose near-zero mean deviation (−0.1 years) did not reflect an absence of pathway-level differences: its youth-like and older-like contributions were of comparable magnitude and offset one another (Fig. 4c). The near-zero total deviation alone could not distinguish this cancellation of large opposing contributions from a tumor closely matching the aging state expected for its tissue of origin. Across the six cancers, tumors therefore occupied composite aging states in which youth-like and older-like contributions coexisted, and their balance determined the tumor’s overall position.

### Breast tumor deviations and their pathway structure replicate across cohorts

The cross-cancer analysis summarized each cancer by its mean contribution profile, leaving open whether these composite states reproduced across cohorts and how their pathway structure varied among tumors within a cancer. We addressed these questions in breast cancer, where PAM50 subtype and *TP53* mutation provide well-established axes of tumor heterogeneity and a large independent cohort is available in SCAN-B (7,744 tumors)^23^. METABRIC provided an additional replication under harmonized processing.^24^ We applied the GTEx-breast clock and its reference unchanged to TCGA-BRCA and SCAN-B, using TCGA-BRCA adjacent-normal tissue as a within-TCGA comparator.

With no retraining, recalibration, or exposure to tumor data, the directly transferred breast clock produced closely aligned youth-like deviations in both TCGA and SCAN-B (Fig. 5a,b). SCAN-B tumors had a mean deviation of −20.2 years, close to the −22.9 years observed in TCGA-BRCA. TCGA-BRCA adjacent-normal tissue showed a smaller deviation of −5.3 years, intermediate between the GTEx-breast reference and the tumor cohorts, consistent with field effects reported in histologically normal tissue adjacent to breast tumors.^25,26^

**Figure 5.**
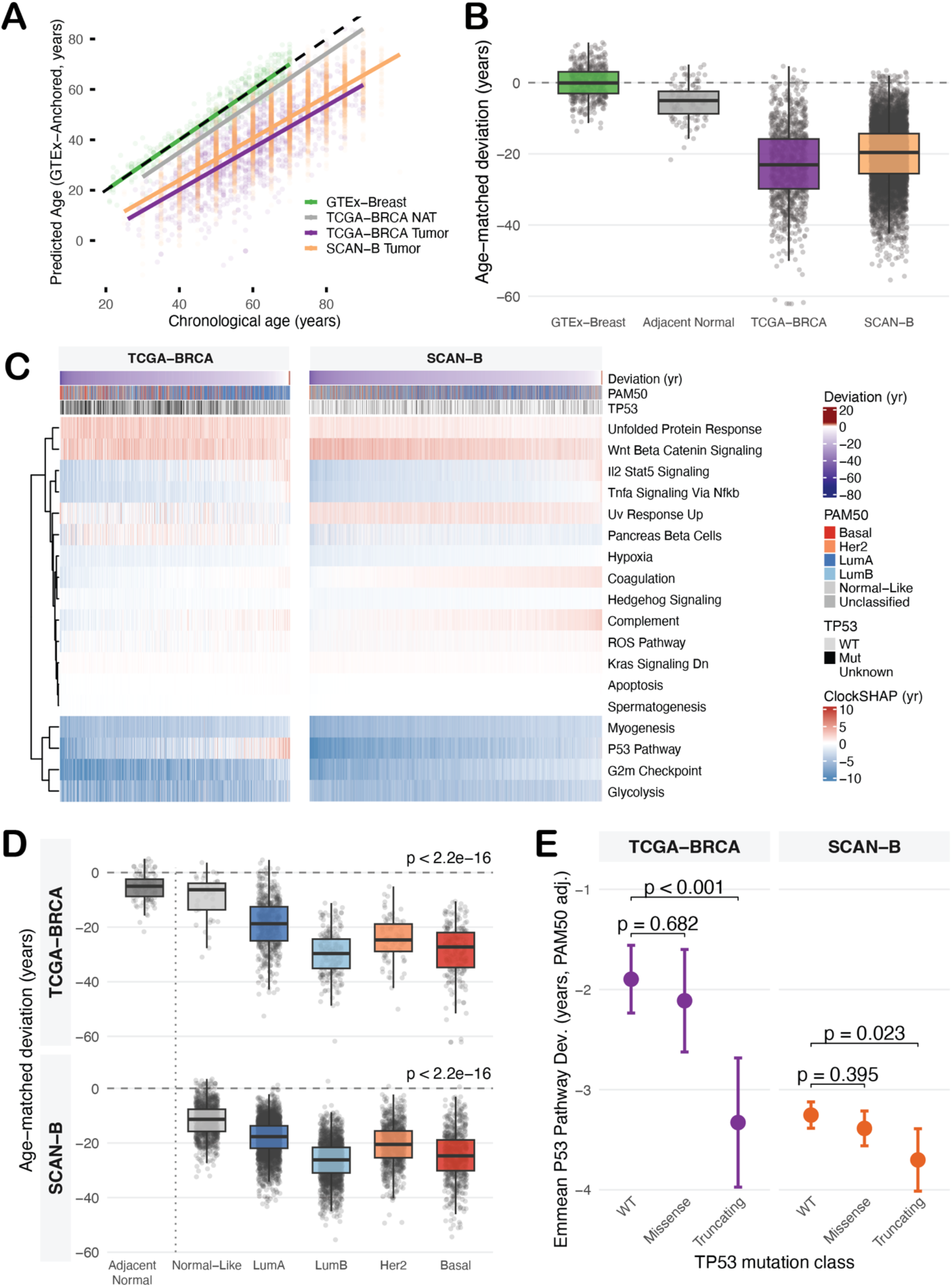
Breast tumor deviations replicate in SCAN-B and structure by subtype and TP53 status. (a) Predicted age (GTEx-anchored as in Fig. 3a) versus chronological age for GTEx breast, TCGA-BRCA adjacent normal, TCGA-BRCA tumors, and SCAN-B tumors; tumors of both cohorts fall well below the identity line. (b) Age-matched deviation by cohort (GTEx breast, n = 482; adjacent normal, n = 112; TCGA-BRCA, n = 1,117; SCAN-B, n = 7,744), with adjacent normal near zero and both tumor cohorts strongly youth-like; boxes show the median and interquartile range, whiskers extend to 1.5× the interquartile range, and points are individual samples. (c) ClockSHAP pathway contributions for TCGA-BRCA and SCAN-B tumors, each column a tumor ordered by deviation and annotated by deviation, PAM50 subtype, and TP53 status; pathway rows are hierarchically clustered and the resulting structure is concordant between cohorts. (d) Age-matched deviation across PAM50 subtypes in each cohort (n = 1,117 TCGA-BRCA and 7,744 SCAN-B tumors), with adjacent normal shown as a reference anchor (excluded from the test); box elements as in b. Subtypes were compared by Kruskal-Wallis (TCGA-BRCA and SCAN-B p < 2.2 × 10⁻¹⁶). (e) PAM50-adjusted P53-pathway ClockSHAP contribution by TP53 mutation class (wild-type, missense, truncating) in patients with TP53 calls (n = 1,080 TCGA-BRCA and 2,284 SCAN-B), modeled within each cohort; points are estimated marginal means averaged over PAM50 subtypes and error bars are 95% confidence intervals, with each class contrasted against wild-type (Dunnett-adjusted).

This agreement extended to the pathway structure driving the deviations (Fig. 5c). Ordering tumors by deviation, with the most youth-like on the left, and decomposing each with ClockSHAP produced closely matching organization in TCGA-BRCA and SCAN-B: G2M checkpoint, glycolysis, and myogenesis contributed youth-like across nearly all tumors, whereas unfolded protein response and Wnt/β-catenin signaling contributed older-like, at comparable magnitudes in both cohorts. The composite state observed at the cancer level was therefore present across individual tumors and reproduced between cohorts.

Beyond this shared pathway architecture, the cohorts also reproduced the structure of tumor-to-tumor variation that was obscured by the cancer-level mean contributions in Fig. 4. Several pathways spanned continuous gradients across tumors rather than contributing uniformly youth-like or older-like, including TNFα–NFκB signaling, IL2–STAT5 signaling, and P53 pathway. The immune-pathway gradients closely matched between TCGA-BRCA and SCAN-B, whereas the P53-pathway contribution differed in range: it spanned both older-like and youth-like values in TCGA-BRCA but remained predominantly youth-like in SCAN-B (Fig. 5c).

The youth-like tumor shift and directions of the major pathway contributions were also recovered in METABRIC using a harmonized analysis in which the GTEx-breast clock was retrained on the gene universe shared across cohorts (Supplementary Fig. 4a,b,e).

Several complementary analyses further supported the validity and stability of the transferred clock. Across the 50 Hallmark scores, GTEx breast and TCGA-BRCA adjacent-normal samples overlapped closely, whereas tumors were displaced from both, arguing that the tumor shift was not simply a GTEx–TCGA cohort effect (Supplementary Fig. 5a). The observed mean BRCA deviation lay far beyond the distribution obtained from age-permuted GTEx clock refits, showing that the shift depended on age-associated structure learned from the normal reference (Supplementary Fig. 5b). Finally, the dominant ClockSHAP pathway pattern was preserved across nearby clock hyperparameter settings, while bootstrap refits consistently selected the major pathways and retained the direction and magnitude of their coefficients, indicating that the decomposition did not depend on a single fitted model (Supplementary Fig. 5c,d).

Finally, we asked whether the youth-like deviation is carried by the tumor cells themselves. In a single-cell breast tumor atlas^27^, pseudobulk profiles built from each donor’s tumor (epithelial) cells were more youth-like than profiles from the same donor’s non-tumor cells in all 64 donors, and each sample’s deviation became more youth-like with increasing tumor-cell fraction, at a rate comparable to that estimated from bulk tumor purity (Supplementary Fig. 6a,b), indicating that the deviation arises in the tumor cells and is diluted, not produced, by the surrounding tissue.

### Breast deviations align with molecular subtype and TP53 mutation class

We next asked whether these replicated deviations aligned with established breast-cancer biology. Although the clock was trained only on normal tissue and received no subtype information, age-matched deviation separated PAM50 subtypes in the same ordered pattern across TCGA-BRCA and SCAN-B (Fig. 5d; Kruskal–Wallis P < 2.2 × 10⁻¹⁶ in both). Normal-like tumors were least youth-like, Luminal A and HER2-enriched tumors occupied intermediate positions, and Luminal B and Basal-like tumors were most youth-like. This ordering tracked established differences in subtype biology and clinical behavior^28^, linking greater youth-like deviation to more aggressive breast tumor states. The clock therefore recovered a major axis of breast-cancer heterogeneity across independently assembled cohorts. A similar subtype ordering was reproduced in METABRIC, with Normal-like tumors least youth-like and Luminal B tumors most youth-like (Supplementary Fig. 4c). The subtype signal was also not attributable to tumor content: in TCGA-BRCA the ordering was preserved after adjustment for tumor purity (Supplementary Fig. 2b), and in the single-cell atlas the subtypes separated among tumor (epithelial) cells alone (Supplementary Fig. 6c).

We next tested whether an individual pathway contribution tracked the biology it represented. We focused on P53 pathway, whose contribution varied widely across tumors, from youth-like to older-like values (−21.7 to +10.2 years in TCGA-BRCA; Fig. 5c). Because P53 pathway carries a positive weight in the breast clock (Supplementary Table 5), lower P53 pathway scores produce more youth-like contributions. The decomposition therefore makes a directional prediction: truncating *TP53* mutations, the clearest loss-of-function class^29^, should be associated with more youth-like P53-pathway contributions. Among patients with *TP53* mutation calls available (n = 1,080 in TCGA-BRCA and n = 2,284 in SCAN-B), we classified mutations as truncating, missense, or wild type and tested their association with the P53-pathway contribution within each cohort, adjusting for PAM50 subtype. Truncating mutations had significantly more youth-like P53-pathway contributions than wild-type tumors in both cohorts (P < 0.001 in TCGA-BRCA and P = 0.02 in SCAN-B; Fig. 5e), whereas missense mutations did not significantly differ from wild type in either, and the truncating association persisted after adjustment for tumor purity in TCGA-BRCA (Supplementary Fig. 2c). In SCAN-B, *TP53* mutations were called from RNA-seq rather than exome sequencing^30^, potentially reducing sensitivity to truncating variants because transcripts containing premature termination codons can be depleted by nonsense-mediated decay.^29^

Together, these results showed that total deviation captured a reproducible axis of PAM50 subtype variation, while ClockSHAP isolated a P53-pathway component associated specifically with truncating *TP53* mutations after adjustment for subtype. These biologically ordered patterns demonstrate that the clock-derived aging states were not arbitrary outputs but interpretable representations of breast-cancer biology.

### Age-matched deviation associates with breast cancer survival

Deviation was associated with overall survival in BRCA (Fig. 4b), but it also separated PAM50 subtypes (Fig. 5d), which differ substantially in clinical behavior.^28^ We therefore asked in TCGA-BRCA and SCAN-B whether the survival association primarily reflected subtype composition or whether deviation retained prognostic information within subtype.

We first confirmed the overall association. Across all tumors, more youth-like deviation was associated with shorter overall survival in both TCGA-BRCA and SCAN-B in Cox models adjusted for chronological age (Fig. 6a), and tumors in the most youth-like deviation tertile had the poorest survival in each cohort (Fig. 6b; log-rank *P* = 0.027 in TCGA-BRCA and *P* < 0.001 in SCAN-B). Thus, both the continuous and tertile-based analyses reproduced the same relationship across two independent cohorts: more youth-like deviation was associated with poorer survival.

**Figure 6.**
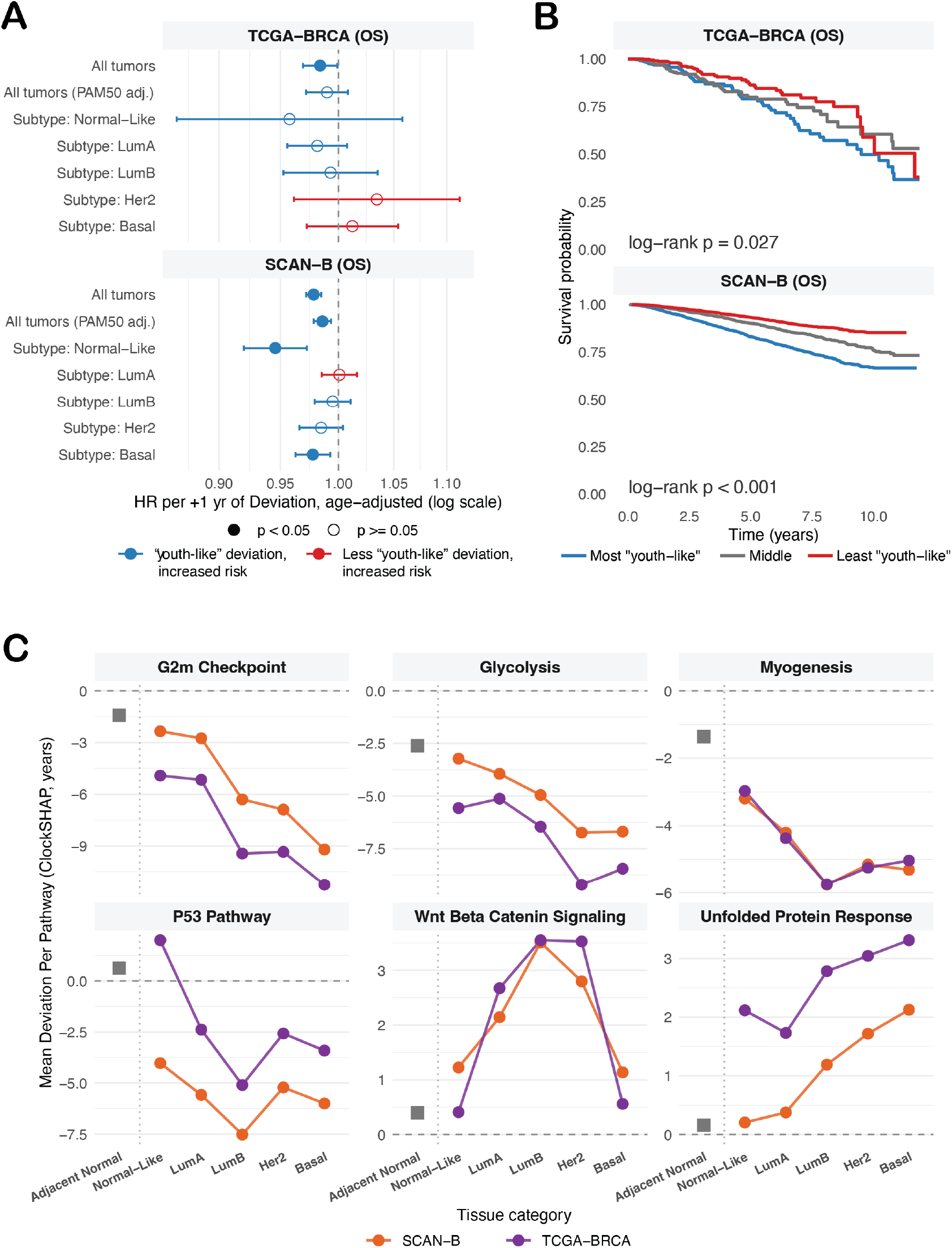
Age-matched deviation is prognostic in breast cancer and its pathway structure tracks subtype. Overall survival analysis in TCGA-BRCA (n = 1,116, 151 events) and SCAN-B (n = 7,742, 1,283 events). (a) Age-adjusted Cox hazard ratio for overall survival per additional year of deviation, across all tumors, adjusted for PAM50 subtype, and within each subtype, for each cohort; error bars are 95% confidence intervals on a log scale, point color indicates the direction of risk (blue, more youth-like deviation associates with higher risk; red, less youth-like with higher risk), and filled points mark p < 0.05 (open, p ≥ 0.05). (b) Kaplan-Meier overall survival by within-cohort deviation tertile (T1, most youth-like; T3, least youth-like), administratively censored at SCAN-B’s maximum follow-up (11.87 years) so both cohorts span the same period; unadjusted log-rank p = 0.027 (TCGA-BRCA) and p < 0.001 (SCAN-B). (c) Mean per-pathway ClockSHAP contribution across PAM50 subtypes for six pathways in each cohort, with TCGA adjacent normal anchored at left; subtype trends are concordant between cohorts.

Part of this association reflected molecular subtype. Adjusting for PAM50 attenuated the deviation effect, which became non-significant in TCGA-BRCA but remained significant in SCAN-B (Fig. 6a). Within individual subtypes, the association was largely absent in TCGA-BRCA (151 events), whereas in SCAN-B (1,283 events) more youth-like deviation remained associated with shorter survival in the Normal-like and Basal-like subtypes (Fig. 6a). METABRIC showed the same attenuation under PAM50 adjustment, and its within-subtype effects were directionally consistent with SCAN-B in Normal-like and Basal-like tumors (Supplementary Fig. 4d), and in TCGA-BRCA the attenuation and the within-subtype estimates were unchanged with tumor purity in the models (Supplementary Fig. 2d). Thus, more youth-like deviation marked more aggressive breast-cancer states partly through subtype composition, while retaining prognostic information within selected subtypes.

### Breast subtypes occupy distinct composite aging states

Given the strong association between deviation and PAM50 subtype, we next asked whether subtypes differed only in the magnitude of their total deviation or also in the pathway composition of that state. ClockSHAP showed that they differed in both (Fig. 6c). G2M checkpoint displayed the clearest youth-like progression, shifting from −2.3 years in Normal-like tumors to −9.2 years in Basal-like tumors in SCAN-B and from −4.9 to −11.3 years in TCGA-BRCA. Unfolded protein response moved broadly in the opposite direction, increasing from +0.2 to +2.1 years in SCAN-B and from +2.1 to +3.3 years in TCGA-BRCA. Glycolysis and myogenesis were likewise more youth-like in Luminal B, HER2-enriched, and Basal-like tumors than in Normal-like tumors. TCGA-BRCA adjacent-normal samples remained closest to zero across all six pathways, indicating that they most closely resembled normal breast tissue.

Other pathways followed subtype-specific patterns. Wnt/β-catenin signaling became increasingly older-like from Normal-like through the luminal and HER2-enriched subtypes before dropping sharply toward zero in Basal-like tumors in both cohorts (Fig. 6c), consistent with relatively higher Wnt/β-catenin activity in Basal-like than luminal breast tumors.^31,32^ The P53-pathway contribution became most youth-like in Luminal B tumors in both cohorts, differing only in the Normal-like group, where it was older-like in TCGA-BRCA (+2.0 years) and youth-like in SCAN-B (−4.0 years). Therefore, the broader cross-cohort difference observed in Fig. 5c was largely concentrated in the Normal-like subtype.

These pathway-by-subtype patterns were also recovered in METABRIC under the harmonized common-gene-universe analysis. The same six pathways showed closely aligned directions, magnitudes, and subtype patterns across TCGA-BRCA, SCAN-B, and METABRIC (Supplementary Fig. 4e).

Breast subtypes therefore occupied distinct composite aging states rather than differing only in how far they sat from their normal-tissue expectation. More youth-like subtypes combined increasingly youth-like proliferative and metabolic contributions with contributions from other programs that were unchanged or increasingly older-like. The composite aging states identified across cancers were also present within breast cancer, where their pathway composition varied systematically by molecular subtype and reproduced across independent cohorts.

## Discussion

Aging and cancer are linked by biological programs that can be shared, divergent, or context dependent. This complexity has been framed through the “meta-hallmarks” of aging and cancer, which characterize their links as equivalent, antagonistic, or ambivalent.^2^ Here, by comparing tumors to the aging expected for their tissue of origin and decomposing their deviations with ClockSHAP, we provide a tissue-resolved quantitative view of these relationships within individual tumors.

Our findings suggest that cancers reshape normal aging programs rather than simply advancing or reversing them. Across 33 cancer types and six molecular layers, age-associated tumor changes retained the tissue-of-origin structure seen in normal tissues, arguing against a single pan-cancer aging axis and motivating tissue-specific comparisons. Within the transcriptomic layer, tumors departed from their tissue-of-origin reference in program-specific directions, with proliferative programs youth-like in nearly every cancer and damage-response and signaling programs often older-like. LGG made this especially clear, since its near-zero mean deviation reflected large opposing contributions rather than close agreement with normal brain aging. Whether this pattern solely reflects what the clocks learned from normal tissue, where proliferative activity declines and damage-response markers accumulate with age, or genuine biological overlap between tumor states and tissue aging cannot be resolved from these data alone and will require comparison across clocks trained on different feature spaces and experimental perturbation of individual pathway contributions.

These results suggest that molecular age is not necessarily a single property of a tumor but a scalar summary of a composite aging state, built from competing programs whose balance depends on the tissue, the cancer, and the features represented by the clock. This may help explain why different clocks or feature spaces have positioned the same cancers as younger, older, or largely unchanged.^8,10–15^ It also suggests that the processes proposed to underlie clock behavior in cancer, including epigenetic maintenance, epigenetic drift, senescence, and stemness, need not be alternatives.^8,10,11^ Each may contribute, with the balance set by which programs a given clock captures.

Breast cancer demonstrated that these aging states track established disease biology. Applied without retraining, a clock trained only on normal breast tissue produced closely matching deviations and pathway contributions across independently assembled tumor cohorts, separated PAM50 subtypes^28^ without receiving subtype information, and linked the P53-pathway contribution specifically to truncating *TP53* mutations.^29^ That a model with no exposure to tumor data, applied to different cohorts on a different continent under different clinical protocols, recovered the same hierarchical structure of pathway contributions is not merely a reproducibility check – it argues that the aging deviations reflect genuine biological signal embedded in the tissue-of-origin aging program, rather than cohort-specific technical or demographic variation. Subtypes also reached their total deviations through different combinations of pathway contributions, showing that the clock resolved distinct subtype-specific states rather than ranking tumors along a single young-to-old axis. That the output was organized by subtype suggests it reflects how cancer biology shapes the tumor’s departure from normal aging, rather than a chronological aging signal independent of tumor state.

Taken together, these results argue that aging-clock studies in disease should define deviations relative to an appropriate normal reference rather than chronological age alone, and that a scalar deviation can remain biologically ambiguous without knowing which features produced it. Age-matched deviation and ClockSHAP provide this reference-based and feature-resolved view. Deviation generalizes conventional *relative age acceleration* (RAA)^9^ by carrying a designated normal reference into new cohorts and is equivalent to RAA within that reference cohort. ClockSHAP decomposes any linear-additive clock exactly into feature-level contributions, distinguishing close agreement with a reference from cancellation among large opposing effects. The approach is not specific to cancer, transcriptomics, or Hallmark pathways. While ClockSHAP faithfully explains what a clock predicts, biological interpretation must still be supported by replication, stability, and concordance with external biology, as done here. We provide ClockSHAP as an R package to support such analyses.

Our study has limitations. The pathway-based clocks predicted chronological age only modestly in held-out normal tissue, reflecting the deliberate use of 50 interpretable features rather than a genome-wide model. Normal references were defined at the organ level and may obscure differences among cell types within the same tissue, as illustrated by LUAD and LUSC sharing a normal lung reference. Bulk transcriptomic profiles also combine cell-intrinsic changes with shifts in tissue composition, although the subtype, survival, and TP53 associations reported here were preserved after adjustment for tumor purity in TCGA-BRCA, and the deviation was carried by the tumor (epithelial) cells rather than the surrounding tissue in a single-cell breast tumor atlas. The use of Hallmark pathways restricted the clocks to broad programs and may omit other aspects of aging biology. Clock analyses were restricted to six cancers with sufficient normal and tumor samples and clear tissue correspondence. Predictions beyond the age range of the training data require particular caution. Broader application would benefit from larger normal-tissue cohorts, finer cellular references informed by single-cell data, and evaluation across additional molecular clocks and disease settings.

Together, these findings support interpreting aging-clock outputs in cancer as tissue-relative composite states rather than literal estimates of tumor age. By grounding deviations in a normal-tissue reference and resolving them into biological programs, the approach opens a path toward understanding how diverse disease states, not only cancers, reshape the aging programs of their tissues of origin.

## Methods

### TCGA data acquisition

To map age-associated molecular change across TCGA primary tumors, six molecular layers were obtained from the NCI Genomic Data Commons (GDC) using TCGAbiolinks^33^ with TCGA-LAML queried without a tumor-type restriction given its blood-derived samples. DNA methylation used Level 3 beta values from the Illumina HumanMethylation450 array (33 cancer types), produced by the GDC SeSAMe pipeline.^34^ RNA-seq used STAR-aligned^35^, unstranded gene-level counts on GRCh38 (33 cancer types). miRNA used BCGSC miRNA read counts (33 cancer types). RPPA used Level 4, replicate-base normalized (RBN) protein expression^36^ for the 32 cancer types with available data (all except TCGA-LAML). Somatic mutation used GDC MAF calls reduced to per-gene binary status (33 cancer types), and somatic copy-number alteration used GISTIC gene-level calls^37^ reduced to binary amplification and deletion status (32 cancer types).

### Sample metadata and covariate specification

Clinical and demographic variables were obtained from the GDC with TCGAbiolinks^33^ and harmonized across layers. Chronological age was defined as age at diagnosis in years. Sex and race were taken from the GDC clinical records, molecular subtype from the TCGA PanCancerAtlas subtype calls^18^ (the consolidated selected subtype, PAM50 for breast cancer^28^), and tumor purity from the TCGA PanCancerAtlas ABSOLUTE estimates.^38^ HPV status for head and neck squamous cell carcinoma was taken from the GDC biospecimen records, with indeterminate calls set to missing. For each cancer and layer, pre-specified rules set the covariates entering the age model in Figure 1: sex when each sex had at least 5 samples and neither exceeded 90% of the cohort; race when at least two categories (excluding “not reported”) each had at least 5 samples, with the analysis then restricted to those categories; tumor purity when at least 75% of samples had a purity estimate; molecular subtype for breast cancer; and tumor grade for cancers with recorded grade. For head and neck squamous cell carcinoma, HPV-positive tumors were excluded so that associations were estimated within HPV-negative disease. A cancer-layer combination was analyzed only if at least 50 samples remained after removing samples missing age or any included covariate.

### Age-association analyses

Within each cancer type, molecular features were tested for association with chronological age as a continuous term, using the pre-specified covariates for that cancer; for the binary somatic mutation and copy-number layers, age was instead the response, regressed on alteration status. Unless otherwise noted, a feature was called age-associated at a Benjamini-Hochberg adjusted p value of 0.05 or below, with direction given by the sign of the age effect. Layer-specific procedures follow.

1. *RNA-seq*. Genes were filtered to non-zero CPM in at least 10 samples, and duplicate gene symbols were resolved by retaining the Ensembl identifier with the highest expression variance. Age associations were estimated with DESeq2^39^ with independent filtering enabled.
2. *DNA methylation.* Probes were removed if entirely missing, non-CpG (“ch”), overlapping common SNPs or SNPs with minor allele frequency above 1%, flagged by the general SeSAMe mask^40^, more than 20% missing, cross-reactive^41^, or on a sex chromosome. Methylation was modeled on M-values (logit-transformed beta) with limma^42^, and summarized at the region level with DMRcate^43^ using lambda = 500 and C = 5 (recommended for Illumina arrays^44^) on hg19 coordinates; regions were called at a Fisher-combined p value of 0.05 or below, with direction given by the sign of the mean methylation difference.
3. *miRNA*. Age associations were estimated with DESeq2^39^ on raw BCGSC miRNA counts, with independent filtering enabled.
4. *RPPA*. Proteins and samples with more than 20% missing values were removed, and RBN values were analyzed without further normalization or imputation; protein-level age associations were estimated with limma.^42^
5. *SCNA*. GISTIC gene-level calls were reduced to per-gene binary amplification and binary deletion status, and age was regressed on each separately (genes with at least 10 samples per group tested); significant genes from both tests were combined.
6. *Somatic mutation.* Per-gene mutation status was taken as the union across mutation categories (nonsense, missense, frameshift, splice-site, translation start site), and age was regressed on mutation status (genes with at least 10 samples per group tested).

### Pan-cancer feature counts (Fig 1A)

For each cancer and layer, features significantly increasing and decreasing with age were tallied by the sign of the age effect and combined across all six layers, with each layer contributing at its native resolution (regions for methylation, genes for mutation and SCNA, individual features otherwise).

### Tissue-of-origin clustering (Fig 1B-C)

Cancers were clustered by the similarity of their per-feature age-effect profiles. For each layer, a feature-by-cancer matrix of per-feature age statistics (moderated t for limma layers, Wald for DESeq2 layers, regression statistic for SCNA) was assembled from features age-associated in at least one cancer, with methylation capped to the 100,000 CpGs most often significant across cancers for tractability. Cancer-by-cancer Pearson similarity was computed over shared features. Similarities estimated from fewer than 100 shared features were discarded, as correlations over a handful of features are unstable. Clustering used methylation, RNA-seq, miRNA, and SCNA. RPPA was excluded because its panel of a few hundred antibodies is too narrow to contribute on equal footing with the genome-wide layers, and somatic mutation because age-associated mutations are too sparse and largely tissue-specific, a median of about one shared feature per cancer pair. Each TCGA cancer type was assigned to an organ system based on the anatomical site of the primary tumor, defined a priori and independent of any clustering result (assignments shown in Fig. 1b).

For the within- versus between-organ comparison (Fig 1C), each layer’s pairwise distances were split into within-organ and cross-organ pairs and restricted to organ systems with at least two cancers. Each layer was evaluated on the cancers assayed in that layer, with the cancer set and the retained organ systems defined after the 100-feature floor was applied. THCA was excluded from the SCNA comparison because all of its pairs fell below the floor. Because each cancer contributes to many pairs, the pairs are not independent. Significance was therefore assessed by permutation: organ-system labels were shuffled across that layer’s cancers and the test statistic, the difference in mean ranks between cross-organ and within-organ pairs, was recomputed over 10,000 permutations. Reported P values are two-sided empirical values from this null.

Clustering was restricted to cancers with results in both methylation and RNA-seq, the two densest layers (CHOL, DLBC, and OV lacked results in one of these and were not clustered). Per-layer similarities were averaged within each cancer pair, counting a layer only where that pair shared at least 100 features, and converted to a distance of 1 minus similarity. Cancers were then clustered with Ward.D2 linkage and optimal leaf ordering.^45^

### recount3 data acquisition

The aging-trend and clock analyses use a uniformly reprocessed RNA-seq resource rather than the multi-layer GDC data above, so that GTEx and TCGA are directly comparable. GTEx and TCGA RNA-seq were obtained from recount3^46^ and scaled to read counts, with genes restricted to protein-coding biotypes, dropping those without a gene symbol or annotated as pseudoautosomal Y duplicates (_PAR_Y). Chronological age was the exact subject age in years (GTEx, from the protected v10 subject phenotype file; TCGA, age at diagnosis from recount3 metadata), and sex was harmonized to a common encoding.

### GTEx-TCGA comparison processing

For the GTEx-TCGA aging-trend comparison, TCGA was restricted to primary disease samples (Primary Tumor, plus Primary Blood Derived Cancer for TCGA-LAML), and all GTEx samples were retained as the normal-tissue reference. Counts for duplicate gene symbols were summed. Projects with fewer than 50 samples with non-missing age were excluded. Age and sex were the only covariates carried forward, applied symmetrically to GTEx and TCGA; tumor-specific covariates such as purity, grade, and stage were intentionally omitted, and compositional differences were instead addressed through cell-type deconvolution.

### Transcriptional age associations and pathway enrichment

Within each GTEx tissue and TCGA cancer, gene-level age associations were estimated with DESeq2^39^ on a gene set shared across all projects within that cohort, modeling chronological age as a continuous term and adjusting for sex when both sexes had at least five samples. A light expression filter (total count of at least 10) was applied, and the age coefficient was extracted with independent filtering disabled so that every gene retained a statistic for meta-analysis and preranked enrichment. Per-project age effects were pooled separately within GTEx (across tissues) and within TCGA (across cancers) by random-effects meta-analysis (metafor, REML^47^) restricted to genes tested in at least 10 projects, yielding a pooled z-statistic and Benjamini-Hochberg FDR per gene. Hallmark pathway aging was summarized with a two-sided Kolmogorov-Smirnov test (hypeR^48^) against the 50 MSigDB Hallmark gene sets^49^, testing each pathway against the up- and down-ranked gene lists and retaining the more significant direction. For the pan-tissue comparison (Fig 2A), genes were ranked by their meta-analysis z-statistic, with genes meta-tested in both cohorts as the shared background. For the tissue-matched comparison (Fig 2C), each TCGA cancer was paired to its GTEx tissue of origin (breast with BRCA, lung with LUAD and LUSC, brain with GBM and LGG, colon with COAD), and enrichment was run on the per-project DESeq2 results ranked by the gene-level Wald statistic, using the intersection of each pair’s gene universes as background. All 50 Hallmark gene sets were tested. For visualization in Fig. 2a, pathways with FDR < 0.05 in at least one cohort were displayed and grouped into broad functional categories for interpretation. Figure 2c was restricted to pathways in the proliferation, immune, and metabolic categories to focus the tissue-matched comparison.

### Cell-type composition and compositional age effects

To provide compositional context for the gene-level results, cell-type proportions were estimated for every retained GTEx tissue and TCGA cancer with EPIC^50^ on log-CPM expression (edgeR^51^, prior count 1) without expression scaling. Because this reference is derived from blood and tumor-microenvironment data rather than normal solid tissue, these estimates are treated as a broad compositional check on the transcriptional age associations rather than a definitive cell-type characterization. Within each project, compositional age effects were modeled with sccomp^52^, using a composition model of chronological age (adjusting for sex under the same five-sample rule) together with an age term in the variability model, with the posterior approximated by the Pathfinder algorithm^53^ and the age term tested per cell type. Per-project effects were pooled within GTEx and within TCGA by random-effects meta-analysis (metafor, REML) per cell type (cell types present in at least three projects) for the pan-tissue comparison (Fig 2B), with Benjamini-Hochberg correction across cell groups within each cohort. For the tissue-matched comparison (Fig 2C), the per-project sccomp age effects were taken directly for each GTEx tissue and its paired TCGA cancers, using the same pairings as the transcriptional analysis.

### Tissue-of-origin clock feature construction

Each TCGA cancer was paired with its GTEx tissue of origin, retaining TCGA primary tumors and, where at least 10 were available, adjacent normal tissue (Solid Tissue Normal). Within each pair, counts were TMM-normalized^54^ and converted to log2 CPM (edgeR, prior count 1), duplicate gene symbols were averaged, and the GTEx, adjacent-normal, and tumor matrices were reduced to their shared protein-coding gene universe. Hallmark pathway activity was scored by single-sample GSEA (ssGSEA)^55^ using GSVA^56^ (alpha 0.25, without ssGSEA normalization) against the 50 MSigDB Hallmark gene sets^49^, producing the 50-pathway feature matrix used for clock training and application. Pathway-level features make the clock output interpretable in terms of biological programs, and rank-based ssGSEA scores reduce dependence on cohort-specific gene expression scales, supporting cross-cohort application. For breast cancer, PAM50 subtype and TP53 status were attached for downstream stratification.

### Tissue-of-origin aging clock training

For each GTEx tissue, an elastic-net model (glmnet^57^) was trained to predict chronological age from the 50 Hallmark ssGSEA pathway scores. The mixing parameter (α, over 0 to 1) and penalty (λ) were tuned by 10-fold age-stratified cross-validation, minimizing cross-validated mean absolute error and selecting the one-standard-error λ; features were standardized within each fold so that no preprocessing information crossed folds. The final model was refit on all GTEx samples for that tissue and stored in a standardized linear-additive form,

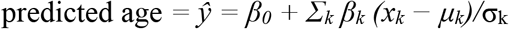

where μ_k_ and σ_k_ are the GTEx training mean and standard deviation of pathway k. This parameterization was verified to reproduce the native glmnet predictions. Equivalently, the clock is linear in the raw pathway scores with per-pathway weight w_k_ = β_k_/σ_k_, the form used by the pathway decomposition below. Each fitted clock was then applied to the TCGA cancers matched to its GTEx tissue, and to adjacent normal tissue where available. Tumor pathway scores were standardized against the GTEx training means and standard deviations, and the clock coefficients were applied without refitting or recalibration, so predicted age for a tumor is the clock’s output on the same feature scale used in training.

### Clock performance evaluation

Clock accuracy was estimated by nested cross-validation, with the full tuning procedure (α grid and one-standard-error λ over 10-fold inner folds) repeated within five age-stratified outer folds so that hyperparameter selection was excluded from performance estimation. Out-of-fold predictions were pooled and summarized as R² (squared Pearson correlation with chronological age), root-mean-square error, and mean absolute error (Supplementary Table 1). The clock deployed for all downstream analyses was refit on the complete GTEx sample, and its fit on those same samples is reported alongside the cross-validated values as training R², root-mean-square error, and mean absolute error. Predicted against chronological age is shown for each cohort and cancer in Supplementary Figure 1.

### ClockSHAP: age-matched deviation and its feature-level decomposition

Because each clock is linear in pathway scores, with per-pathway weight w_k_, its prediction admits an exact decomposition into per-pathway contributions relative to an age-matched normal reference, inspired by the SHAP framework.^58^ We implement this as ClockSHAP (montilab/ClockSHAP).

A reference profile was first fit on the GTEx normal samples: for each Hallmark pathway k, the age-conditioned expected score E[X_k_ | Age] was estimated by regressing the pathway score on chronological age across GTEx (linear), giving the pathway value typical of normal tissue at any age. Evaluating all 50 pathway trends at a given chronological age yields the age-matched reference profile, the vector of expected pathway values for a normal sample of that age. For a tumor sample *i* with pathway scores *x_i,k_* and chronological age *Age_i_*, the age-matched deviation is the clock applied to *x_i_* minus the clock applied to the reference profile at *Age_i_*. Both terms are therefore clock predictions, one on the sample’s observed pathway scores and one on a constructed profile, so the intercept cancels and the deviation reduces to a weighted sum of departures from that reference,

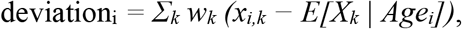

which measures each tumor against the age-matched normal expectation rather than its absolute predicted age. This construction generalizes *relative age acceleration* (RAA), the residual of predicted age regressed on chronological age within the cohort under study. Because the clock is linear in the pathway scores and each reference trend is linear in age, the expected prediction is itself linear in age. When the reference is fit in the cohort being analyzed, it coincides exactly with the ordinary least squares fit of predicted on chronological age, so deviation and RAA are identical sample for sample and deviations are centered at zero by construction. The two diverge when the reference is fit in one cohort and applied to another, as done here: the expectation is the reference cohort’s trend carried forward unchanged, so tumors are measured against normal tissue rather than recentered within the tumor cohort. No post hoc calibration is applied to any prediction.

Each pathway’s exact additive contribution to the deviation is

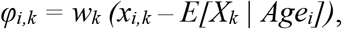

so that *Σ_k_ φ_i,k_* = deviation_i_ exactly. Each *φ_i,k_* is expressed in years and signed, positive where the pathway pushes the tumor toward an older-like prediction and negative toward a younger-like one. For visualization, predicted age is anchored as chronological age plus deviation, so that the age-matched normal reference coincides with the identity line and vertical distance from it equals the deviation (Fig. 3a, 5a). The construction is clock-agnostic and feature-agnostic, applying to any linear clock and any feature set. ClockSHAP was computed for every TCGA tumor, and for adjacent normal samples where available, using the GTEx-fit reference and the matched-tissue clock. The resulting per-sample deviation and *φ* values underlie the cross-cancer deviation landscape, the pathway-decomposition figures, and the survival analysis. For the cross-cancer summaries, each cancer was represented by the mean *φ* per pathway. In Fig. 4c, the four largest-magnitude contributors were shown individually and the remaining pathways aggregated as “Other.” Fig. 4d was restricted to pathways with an absolute mean contribution of at least one year in two or more cancers. ClockSHAP is available as an R package (montilab/ClockSHAP), with documentation providing additional guidance on its use and interpretation.

### Tumor deviation and overall survival

Within each cancer, the association between age-matched deviation and overall survival was tested with a Cox proportional-hazards model, Surv(time, event) ∼ age + deviation, using the age-matched deviation as the continuous predictor. Overall survival was drawn from the TCGA Clinical Data Resource^59^; cancers with fewer than 10 events were not modeled. The identical model was applied across cancers for the cross-cancer comparison (Fig 4B), reporting the hazard ratio per additional year of deviation. The proportional-hazards assumption was assessed with scaled Schoenfeld residuals and held for the deviation term in all cancers except LGG, whose hazard ratio is reported as a time-averaged effect (Supplementary Table 3).

### SCAN-B replication cohort

SCAN-B breast tumor RNA-seq^23,60^ was reduced to one replicate per sample by retaining the most aligned read pairs, and restricted to invasive primary tumors, excluding normal, lymph-node, and in-situ specimens, to match the TCGA-BRCA scope. Expression was processed into the 50-pathway Hallmark ssGSEA feature space as for the training cohorts, and chronological age was taken from SCAN-B’s 5-year binned age variable, which labels each bin by its upper bound. PAM50 subtype and survival endpoints (overall survival, recurrence-free interval, and distant recurrence-free interval) were carried from the SCAN-B annotations, and the breast clock and its GTEx-Breast reference were applied unchanged to obtain predicted age, age-matched deviation, and per-pathway *φ*.

### Breast subtype and TP53 analyses

Within TCGA-BRCA and within SCAN-B, age-matched deviation was compared across PAM50 subtypes by Kruskal-Wallis test, with adjacent normal shown as reference (Fig 5D). For the TP53 analysis (Fig 5E), TP53 status was taken from each cohort’s mutation annotations (TCGA-BRCA from cBioPortal^61^, SCAN-B from Brueffer et al.^30^) and classified identically as truncating (nonsense, frameshift, or splice-site), missense, or wild-type, with in-frame and splice-region variants set aside. The P53-pathway ClockSHAP contribution was modeled on TP53 class adjusting for PAM50 subtype within each cohort, and estimated marginal means per class were contrasted against wild-type with Dunnett adjustment. TCGA-BRCA patients with more than one tumor profile were represented by the mean of their contributions.

### Breast cancer survival

For breast cancer, the age-adjusted deviation model was applied to both TCGA-BRCA and SCAN-B in three forms: across all tumors, additionally adjusted for PAM50 subtype, and separately within each subtype (Fig 6A). Tumors in each cohort were divided into deviation tertiles and compared by Kaplan-Meier overall survival with a log-rank test (Fig 6B). Cox models used all available follow-up, while Kaplan–Meier analyses were administratively censored at 11.87 years in both cohorts to match the maximum follow-up available in SCAN-B.

### Breast clock and decomposition stability

Robustness of the breast clock and its ClockSHAP decomposition was assessed with four checks. First, GTEx normal, adjacent-normal, and tumor samples were projected by principal-component analysis on the standardized Hallmark ssGSEA features, confirming the cohorts share a common feature space rather than separating along a dominant technical axis. Second, a permuted-age null was constructed by shuffling GTEx ages, refitting the clock at the tuned hyperparameters, and recomputing the mean tumor deviation, repeated 1,000 times, against which the observed mean deviation was compared. Third, the clock was refit across a neighborhood of the tuned hyperparameters (the mixing parameter offset by ±0.25 and the penalty scaled by 0.5, 1, and 2), recomputing the mean per-pathway contribution for each. Fourth, the clock was refit on 1,000 bootstrap resamples of the GTEx training samples at the tuned hyperparameters, summarizing each pathway’s coefficient distribution, selection frequency, and sign stability.

### METABRIC validation on a common gene universe

As a further replication, the breast analysis was repeated on a gene universe shared across all cohorts to include METABRIC, a microarray cohort.^24^ METABRIC expression was taken as microarray log-intensity and collapsed to gene symbols by mean, with age at diagnosis and native PAM50 subtype carried from the METABRIC annotations. GTEx, TCGA, SCAN-B, and METABRIC were reduced to the intersection of their gene symbols, Hallmark pathways were scored by ssGSEA on this common universe, and the elastic-net clock was retrained on GTEx using the same procedure as the primary clock. The retrained clock and its GTEx reference were applied to all three tumor cohorts to obtain deviation and per-pathway φ, and the subtype and survival analyses were carried out as before.

### Tumor purity

ABSOLUTE purity estimates (above) were joined to tumor profiles by patient barcode; tumors without an estimate were excluded only from purity analyses, and SCAN-B has no purity estimate. Within each cancer, deviation was related to purity by Spearman correlation and by linear regression on chronological age and purity. For the subtype comparison, each tumor’s deviation was adjusted to the cohort-mean purity using the purity coefficient from a linear model of deviation on PAM50 subtype, chronological age, and purity. Cox and TP53 models were refit with purity as an additional covariate.

### Single-cell atlas analysis

To test directly whether the deviation is carried by tumor cells, we used an integrated single-cell atlas of untreated breast tumors^27^ restricted to donors with a recorded age (72 donors from four of its contributing studies). For each donor, pseudobulk profiles were built by summing raw counts over all cells, over epithelial cells (the tumor compartment), over all non-epithelial cells (the non-tumor compartment), and over epithelial cells flagged as copy-number abnormal by the atlas’s inferCNV calls (a stricter malignant subset). Profiles with fewer than 100 epithelial, 50 non-epithelial, or 100 flagged cells were excluded, leaving 64 donors. Profiles were processed into Hallmark ssGSEA scores and scored with the GTEx-breast clock, its reference, and ClockSHAP exactly as for SCAN-B. Because pseudobulk profiles carry a study-specific platform offset, compartments were compared within donor by Wilcoxon signed-rank test. Cross-donor analyses were repeated with study as a covariate.

## Data Availability

Datasets were obtained from the following sources:

• TCGA molecular and clinical data, NCI Genomic Data Commons: https://portal.gdc.cancer.gov

• GTEx and TCGA RNA-seq counts, recount3: https://rna.recount.bio

• GTEx subject ages (controlled access), dbGaP accession phs000424.v10.p2: https://www.ncbi.nlm.nih.gov/gap/

• SCAN-B gene expression and clinical data, Mendeley Data: https://doi.org/10.17632/yzxtxn4nmd

• SCAN-B TP53 mutation calls, Brueffer et al.^30^

• METABRIC expression, clinical, and TP53 data, cBioPortal: https://www.cbioportal.org/study/summary?id=brca_metabric

• Integrated single-cell breast cancer atlas, CELLxGENE: https://cellxgene.cziscience.com/collections/9432ae97-4803-4b9f-8f64-2b41e42ad3cb

Per-feature age-association results, covariate sets and sample sizes, and Hallmark pathway enrichment results for every analyzed cancer and layer are available in the Pan-Cancer Aging Explorer: https://montilab.github.io/pancancer-aging-explorer/

## Code Availability

Scripts for reproducing results and figures: https://github.com/montilab/bock_et_al_pancancer_aging_clocks

ClockSHAP R package: https://github.com/montilab/ClockSHAP

Pan-Cancer Aging Explorer source: https://github.com/montilab/pancancer-aging-explorer

---

**Supplementary Figure 1.**
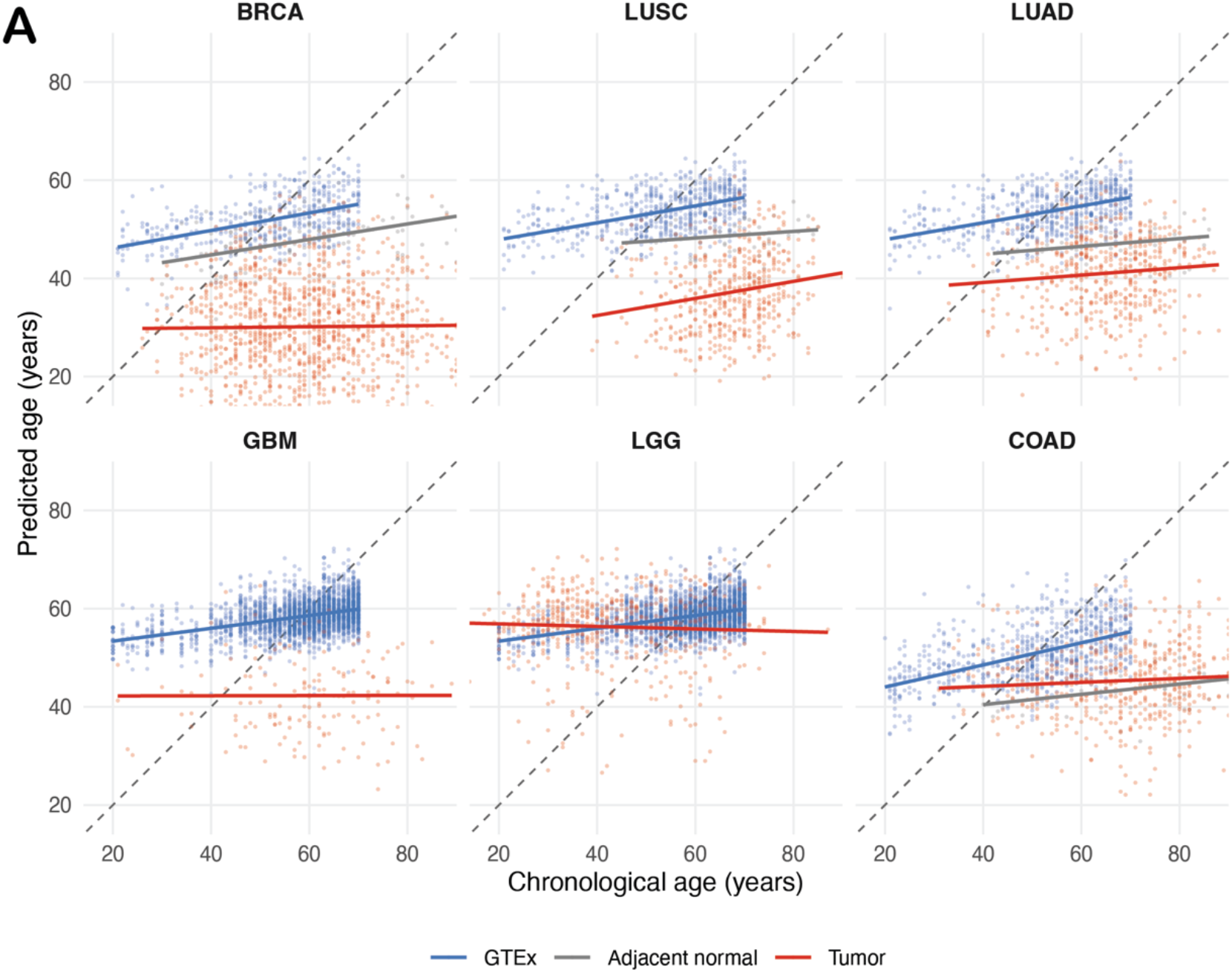
Predicted against chronological age for each cancer. Raw clock output for the GTEx tissue of origin, TCGA adjacent normal tissue, and TCGA primary tumors, one panel per cancer. The dashed line marks equality between predicted and chronological age; solid lines are within-cohort linear fits of predicted on chronological age. The GTEx cohort is the clock’s training set and appears in both panels of each shared tissue, lung in LUAD and LUSC and brain in GBM and LGG. Axes span the chronological age range across all samples and are shared across panels; 57 tumors with predicted ages below 14 years (56 TCGA-BRCA, 1 TCGA-LUSC) fall outside the plotted area and are included in the fitted lines. Cohort sample sizes are given in Supplementary Table 2.

**Supplementary Figure 2.**
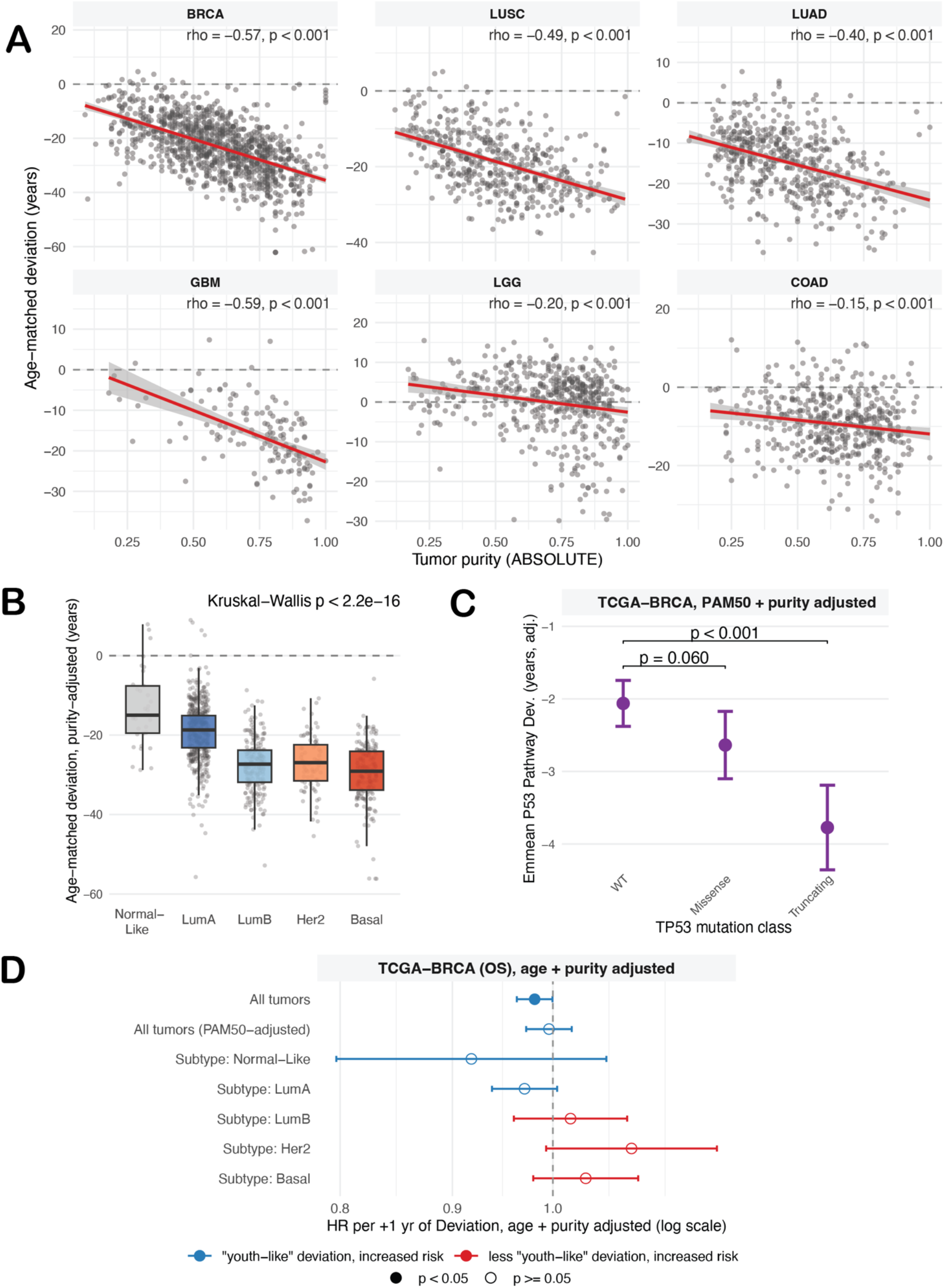
Age-matched deviation and tumor purity. TCGA tumors with an ABSOLUTE purity estimate. (a) Age-matched deviation against tumor purity in each cancer (n = 1,069 BRCA, 483 LUSC, 508 LUAD, 145 GBM, 506 LGG, 484 COAD); lines are linear fits with 95% confidence bands, and Spearman rho and p are given per panel. (b) Purity-adjusted age-matched deviation across PAM50 subtypes in TCGA-BRCA (n = 1,069), drawn as in Figure 5d; Kruskal-Wallis p < 2.2 × 10⁻¹⁶. (c) PAM50- and purity-adjusted P53-pathway contribution by TP53 mutation class in TCGA-BRCA (n = 1,032 patients), drawn as in Figure 5e. (d) Age- and purity-adjusted Cox hazard ratio for overall survival per additional year of deviation in TCGA-BRCA (n = 1,068; 143 events), drawn as in Figure 6a. SCAN-B has no purity estimate.

**Supplementary Figure 3.**
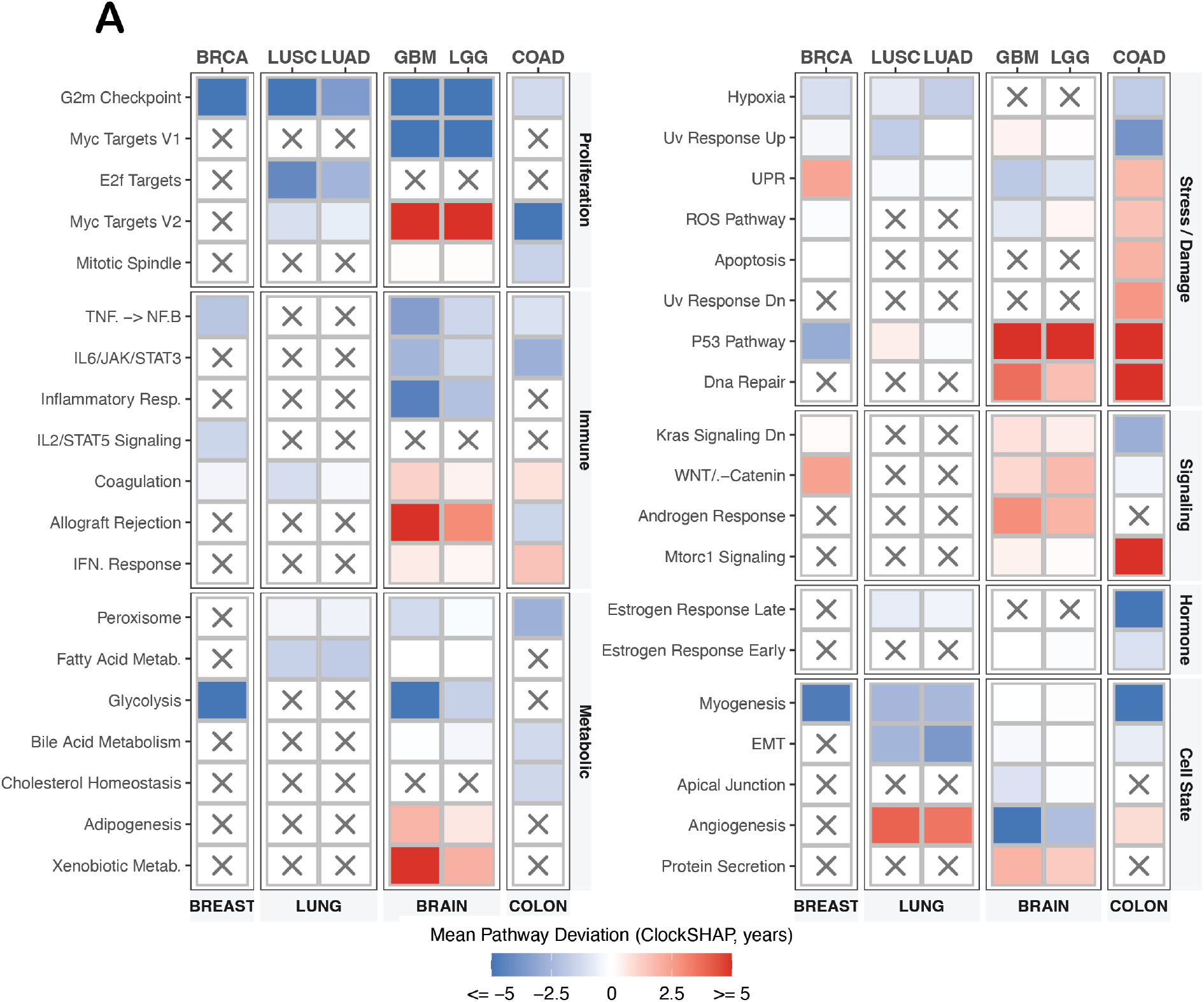
Full cross-cancer pathway deviation landscape. Mean per-pathway age-matched deviation (ClockSHAP) across the six emphasis cancers, extending Figure 4d by relaxing the cross-cancer recurrence requirement. Rows are all pathways whose mean contribution reaches at least one year in magnitude in at least one cancer, grouped by category; columns are cancers grouped by tissue of origin; color is the mean contribution in years (capped at ±5), and a grey cross marks pathways not retained by that cancer’s clock. This view exposes single-cancer pathway signals excluded from Figure 4d, which required recurrence in at least two cancers.

**Supplementary Figure 4.**
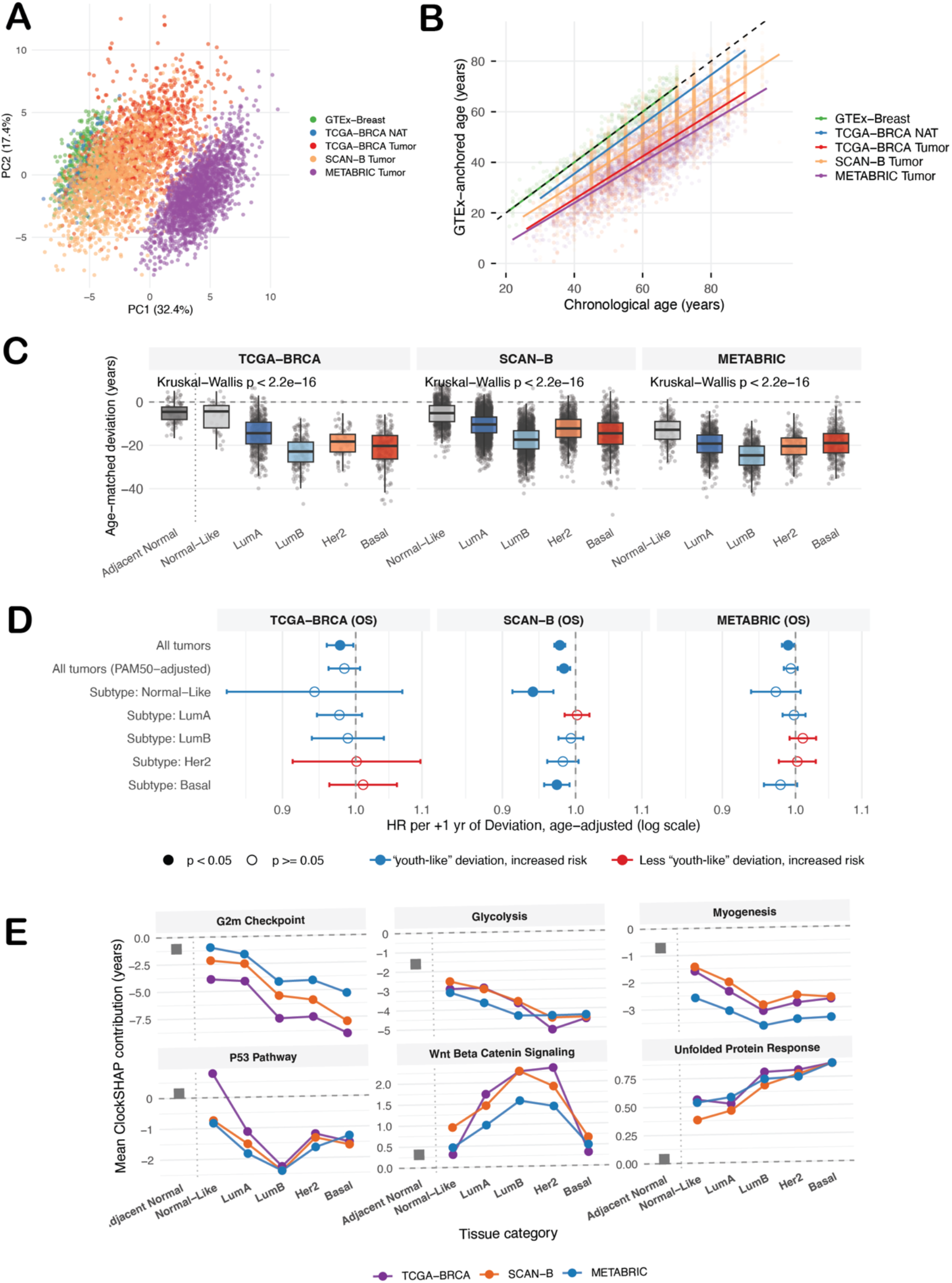
METABRIC validation on a common gene universe. Replication of the breast deviation analysis across GTEx Breast, TCGA-BRCA, SCAN-B, and METABRIC, with all cohorts scored on their shared gene universe and the elastic-net clock retrained on that universe. (a) Principal-component analysis of the Hallmark ssGSEA features across cohorts, with SCAN-B randomly subsampled (to 1,500) (b) Predicted age (GTEx-anchored) versus chronological age across cohorts. (c) Age-matched deviation across PAM50 subtypes in each tumor cohort (n = 1,117 TCGA-BRCA, 7,744 SCAN-B, 1,974 METABRIC tumors), with adjacent normal shown as a reference anchor (excluded from the test); boxes show the median and interquartile range, whiskers extend to 1.5× the interquartile range, and points are individual tumors. Subtypes were compared by Kruskal-Wallis (TCGA-BRCA p = 1.5 × 10⁻⁵⁷; SCAN-B p < 2.2 × 10⁻¹⁶; METABRIC p = 2.4 × 10⁻⁸⁹). (d) Age-adjusted Cox hazard ratio for overall survival per additional year of deviation, across all tumors, adjusted for PAM50, and within each subtype, for each cohort; error bars are 95% confidence intervals on a log scale (color and fill as in Figure 6a). (e) Mean per-pathway ClockSHAP contribution across PAM50 subtypes for six pathways in each cohort, with TCGA adjacent normal anchored at left.

**Supplementary Figure 5.**
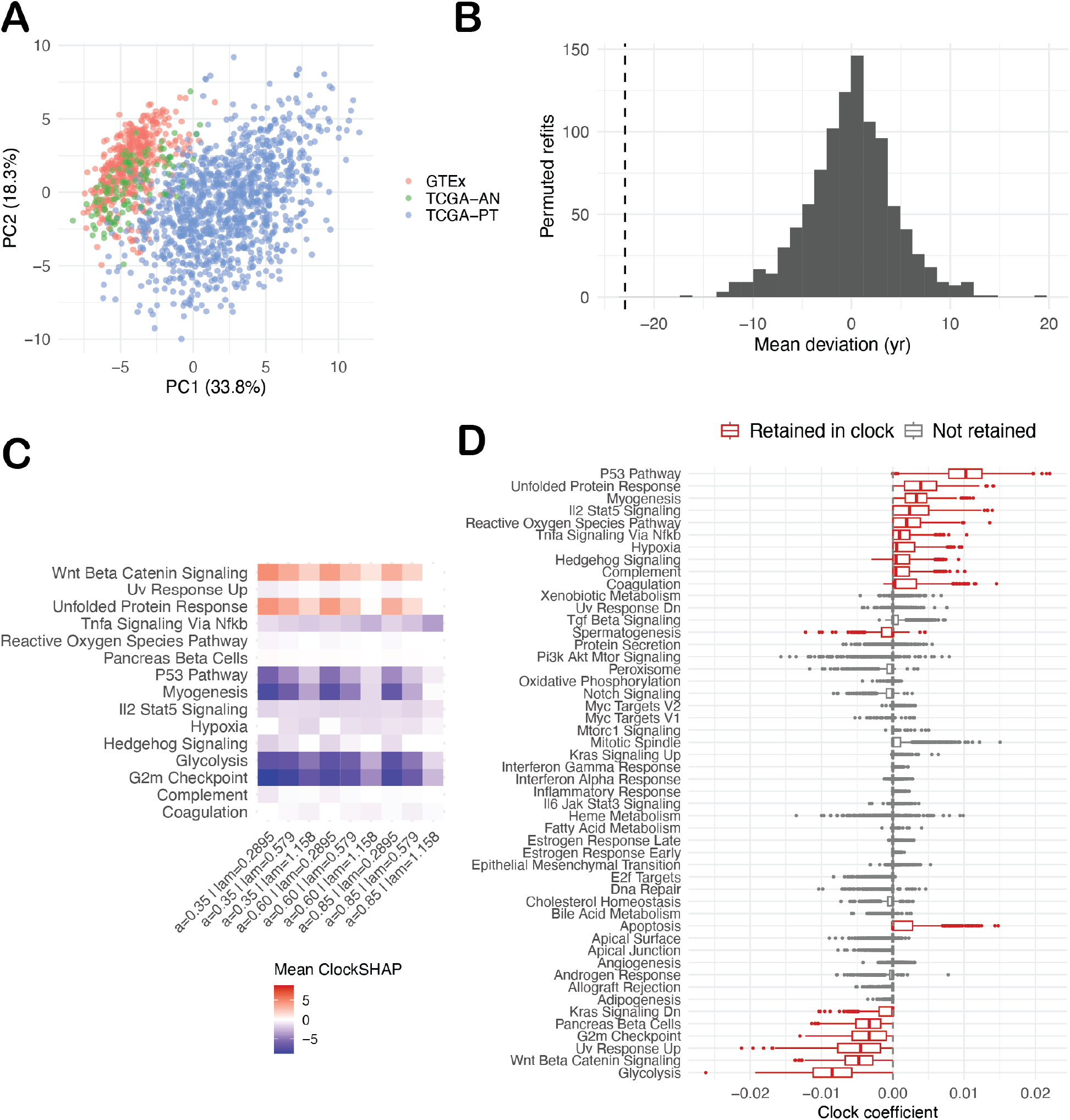
Stability of the breast aging clock and its ClockSHAP decomposition. (a) Principal-component analysis of the standardized Hallmark ssGSEA features for GTEx breast (n = 482), TCGA-BRCA adjacent normal (n = 112), and TCGA-BRCA tumor (n = 1,117) samples, colored by cohort. (b) Permuted-age null: the distribution of mean tumor deviation across 1,000 clock refits on GTEx with shuffled ages, refit at the tuned hyperparameters, with the observed mean deviation marked (dashed line; two-sided empirical P < 0.001). (c) Mean per-pathway ClockSHAP contribution across a neighborhood of the tuned hyperparameters (mixing parameter offset by ±0.25, penalty scaled by 0.5, 1, and 2), showing the pathway structure is stable to tuning. (d) Clock coefficients across 1,000 bootstrap resamples of the GTEx training set refit at the tuned hyperparameters, with pathways retained in the final clock (red) distinguished from those not retained (grey).

**Supplementary Figure 6.**
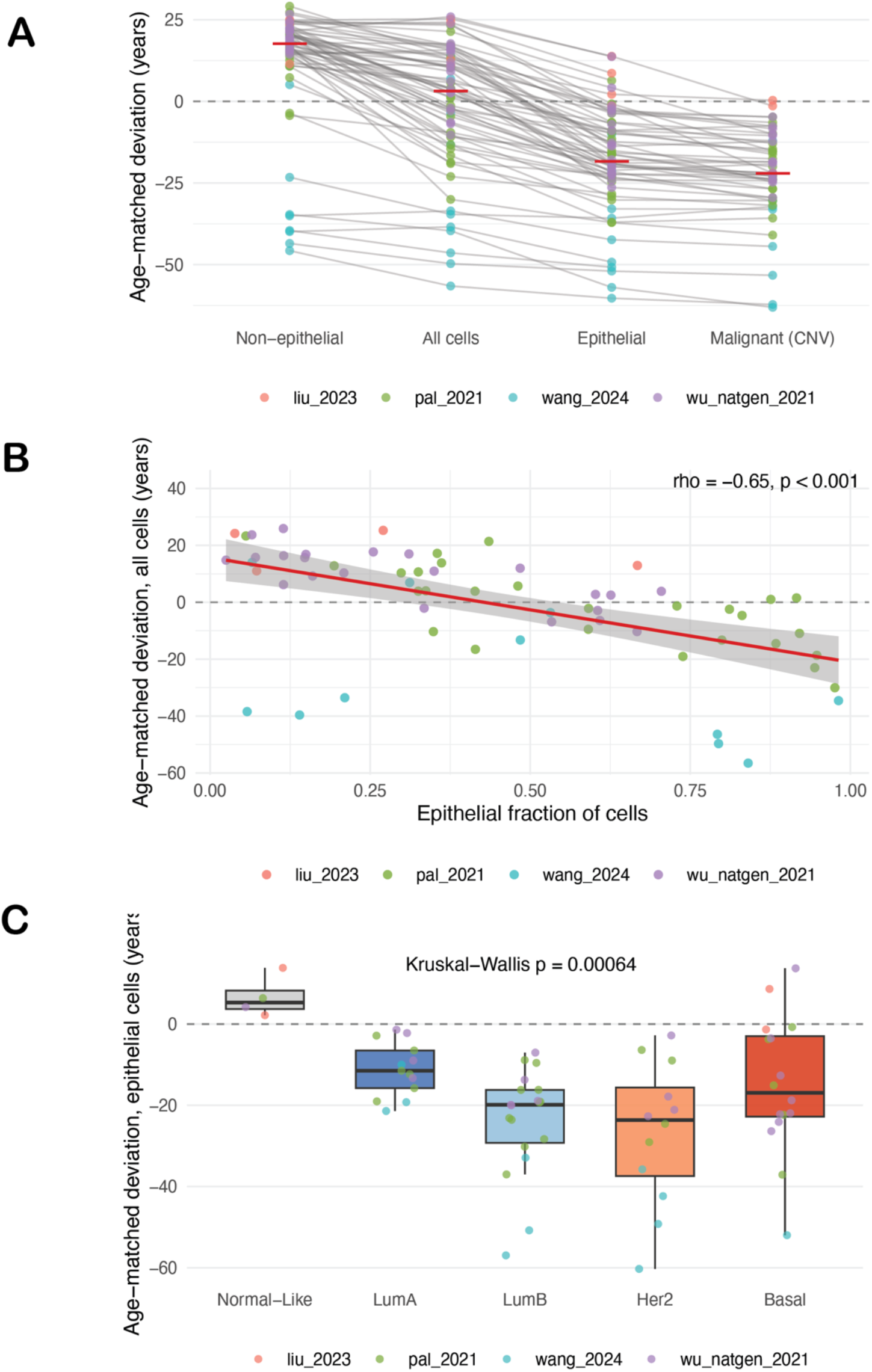
Age-matched deviation by cellular compartment in a single-cell breast cancer atlas. Donor-level pseudobulk profiles from a single-cell atlas of untreated breast tumors (64 donors after cell-count thresholds, four studies), scored with the GTEx-breast clock unchanged (Methods). Tumor (epithelial) cells, non-tumor (all non-epithelial) cells, and a stricter malignant subset of epithelial cells flagged as copy-number abnormal were pseudobulked separately for each donor. Absolute deviations carry a study-specific platform offset and are not interpreted; comparisons within donor cancel it. (a) Deviation of each compartment, one line per donor (n = 64; malignant subset n = 49, donors with at least 100 flagged cells); red bars are medians; points are colored by study. The tumor (epithelial) compartment was more youth-like than the non-tumor compartment in 64 of 64 donors (median −32 years; Wilcoxon signed-rank p < 10⁻¹⁰), and the malignant subset in 49 of 49 (median −36). (b) Deviation of the all-cell profile against the fraction of tumor (epithelial) cells (n = 64; Spearman ρ = −0.65); line, linear fit with 95% confidence band. (c) Deviation of the tumor (epithelial) compartment by donor PAM50 subtype (n = 4, 13, 19, 12, 16); box elements as in Figure 5d; Kruskal-Wallis p = 6 × 10⁻⁴.

**Supplementary Table 1.** GTEx tissue-of-origin clock performance and cohort descriptors. Performance of the four GTEx tissue clocks (breast, brain, lung, colon), reported as both in-sample fit and cross-validated performance. For each clock: the GTEx tissue and its number of training samples; the median and range of chronological age; the number of Hallmark pathways retained with non-zero coefficients in the deployed clock, out of 50 input pathways; training R² and root-mean-square error, from the deployed clock evaluated on the GTEx samples it was fit on; out-of-fold R² and root-mean-square error from nested cross-validation, with the full elastic-net tuning procedure repeated within five age-stratified outer folds; and the alpha and lambda selected by the full-data tuning used for the deployed clock. R² is the squared Pearson correlation between predicted and chronological age in both cases, and root-mean-square error is in years. The clock applied to all tumor analyses was refit on the complete GTEx sample for each tissue.

| **Tissue** | **n** | **Age, median (range)** | **Pathways** | **Training R2 (RMSE)** | **Nested-CV R2 (RMSE)** | **alpha** | **lambda** |
|---|---|---|---|---|---|---|---|
| BREAST | 482 | 54 (21-70) | 18 of 50 | 0.222 (11.6) | 0.136 (12.2) | 0.6 | 0.579 |
| BRAIN | 2931 | 61 (20-70) | 40 of 50 | 0.168 (9.5) | 0.133 (9.7) | 0.2 | 0.1796 |
| LUNG | 655 | 56 (21-70) | 16 of 50 | 0.216 (10.7) | 0.166 (11.0) | 0.6 | 0.579 |
| COLON | 822 | 54 (20-70) | 31 of 50 | 0.262 (11.6) | 0.166 (12.3) | 0.5 | 0.1796 |

**Supplementary Table 2.** Age-matched deviation by cohort. Deviation summary for the GTEx tissue-of-origin reference, TCGA adjacent normal tissue, and TCGA primary tumors across the six cancers. Deviation values are in years relative to the GTEx age-conditioned reference. The GTEx cohort has a mean deviation of zero by construction, since the clock and its reference are fit within it. Reported per cohort are sample count, median and range of chronological age, and the mean, standard deviation, 95% confidence interval of the mean, median, and interquartile range of deviation. The GTEx reference appears once per tissue, shared by LUAD and LUSC in lung and by GBM and LGG in brain. Adjacent normal tissue was carried forward only where at least 10 Solid Tissue Normal samples were available, which excluded GBM and LGG.

| Cohort | n | Age, median (range) | Mean Deviation (SD) | 95% CI | Median (IQR) |
|---|---|---|---|---|---|
| GTEx breast | 482 | 54 (21-70) | -0.0 (4.4) | -0.4 to 0.4 | -0.1 (-3.0, 3.0) |
| BRCA adjacent normal | 112 | 56 (30-90) | -5.3 (5.0) | -6.3 to -4.4 | -5.0 (-8.7, -2.4) |
| BRCA tumor | 1117 | 58 (26-90) | -22.9 (10.4) | -23.5 to -22.3 | -23.1 (-29.8, -15.8) |
| GTEx lung | 655 | 56 (21-70) | -0.0 (3.9) | -0.3 to 0.3 | 0.1 (-2.6, 2.8) |
| LUSC adjacent normal | 49 | 68 (45-85) | -7.4 (3.5) | -8.4 to -6.4 | -7.1 (-10.4, -4.7) |
| LUSC tumor | 491 | 68 (39-90) | -18.8 (8.0) | -19.5 to -18.1 | -19.2 (-24.8, -13.2) |
| LUAD adjacent normal | 59 | 66 (42-86) | -8.8 (3.3) | -9.7 to -8.0 | -8.2 (-10.4, -6.9) |
| LUAD tumor | 520 | 66 (33-88) | -14.6 (7.8) | -15.3 to -13.9 | -14.0 (-20.2, -9.1) |
| GTEx brain | 2931 | 61 (20-70) | 0.0 (3.0) | -0.1 to 0.1 | -0.1 (-1.9, 1.8) |
| GBM adjacent normal | 0 | - | - | - | - |
| GBM tumor | 156 | 60 (21-89) | -16.3 (8.1) | -17.5 to -15.0 | -16.8 (-21.3, -10.8) |
| LGG adjacent normal | 0 | - | - | - | - |
| LGG tumor | 513 | 41 (14-87) | -0.1 (7.9) | -0.7 to 0.6 | 1.5 (-2.9, 4.8) |
| GTEx colon | 822 | 54 (20-70) | 0.0 (5.1) | -0.3 to 0.3 | -0.3 (-3.5, 3.2) |
| COAD adjacent normal | 41 | 74 (40-90) | -11.7 (4.7) | -13.2 to -10.2 | -11.5 (-15.0, -9.6) |
| COAD tumor | 500 | 69 (31-90) | -9.4 (7.7) | -10.1 to -8.7 | -9.2 (-14.3, -4.4) |

**Supplementary Table 3.** Overall-survival Cox results for tumor deviation, with proportional-hazards diagnostics. Per-cancer Cox proportional-hazards models of overall survival on GTEx-anchored age-matched deviation, adjusted for chronological age, reporting the hazard ratio per additional year of deviation with 95% confidence interval, the model p value, and scaled-Schoenfeld proportional-hazards diagnostics for both the deviation term and the global model. The proportional-hazards assumption held for the deviation term in all cancers except LGG (Schoenfeld p = 0.001), whose hazard ratio is therefore a time-averaged effect. In GBM the global test flagged a violation attributable to the chronological-age covariate, while the deviation term itself was proportional and its effect on survival was null.

| Cancer | Tissue | n | Events | HR (95% CI) | p | Schoenfeld p (deviation) | Schoenfeld p (global) | Deviation_PH |
|---|---|---|---|---|---|---|---|---|
| **LGG** | BRAIN | 512 | 125 | 0.951 (0.933-0.969) | 2.31E-07 | 0.00107 | 0.00016 | Violated |
| **BRCA** | BREAST | 1116 | 151 | 0.984 (0.969-0.999) | 0.0358 | 0.279 | 0.549 | Met |
| **LUAD** | LUNG | 520 | 185 | 0.993 (0.975-1.011) | 0.439 | 0.757 | 0.362 | Met |
| **GBM** | BRAIN | 156 | 124 | 1.000 (0.978-1.023) | 0.97 | 0.106 | 0.0306 | Met |
| **LUSC** | LUNG | 488 | 210 | 1.013 (0.996-1.029) | 0.128 | 0.209 | 0.326 | Met |
| **COAD** | COLON | 499 | 109 | 1.039 (1.013-1.066) | 0.00349 | 0.463 | 0.705 | Met |

**Supplementary Table 4.** Overall-survival Cox results for tumor deviation with adjustment for tumor purity. Per-cancer Cox proportional-hazards models of overall survival on GTEx-anchored age-matched deviation in the tumors with an ABSOLUTE purity estimate, adjusted for chronological age alone and for chronological age and purity. Reported per cancer are the number of tumors and events, and for each model the hazard ratio per additional year of deviation with 95% confidence interval and the model p value. Estimates on all eligible tumors are given in Supplementary Table 3.

| **Cancer** | **Tumors (events)** | **HR per year, age-adjusted (95% CI)** | **p** | **HR per year, age + purity adjusted (95% CI)** | **p** |
|---|---|---|---|---|---|
| **BRCA** | 1068 (143) | 0.985 (0.969-1.001) | 0.059 | 0.981 (0.963-1.000) | 0.046 |
| **LUSC** | 480 (207) | 1.014 (0.998-1.031) | 0.09 | 1.016 (0.998-1.035) | 0.083 |
| **LUAD** | 508 (176) | 0.988 (0.970-1.007) | 0.207 | 0.997 (0.977-1.017) | 0.734 |
| **GBM** | 145 (115) | 0.999 (0.977-1.022) | 0.963 | 0.983 (0.957-1.011) | 0.226 |
| **LGG** | 505 (125) | 0.952 (0.934-0.971) | <0.001 | 0.950 (0.932-0.969) | <0.001 |
| **COAD** | 483 (105) | 1.038 (1.011-1.065) | 0.005 | 1.038 (1.012-1.066) | 0.005 |

**Supplementary Table 5. Coefficients of the GTEx tissue-of-origin clocks.** Fitted elastic-net coefficients of the four GTEx tissue clocks (breast, lung, brain, colon), with one row per tissue and Hallmark pathway and one intercept row per tissue. Reported per pathway are the coefficient in years per GTEx standard deviation of the pathway score (βk), the GTEx training mean (μk) and standard deviation (σk) of the pathway score, the per-pathway weight used by ClockSHAP (wk = βk/σk), and whether the pathway was retained with a non-zero coefficient. Predicted age is the intercept plus the sum over pathways of βk(xk − μk)/σk (Methods). The lung clock is shared by LUAD and LUSC and the brain clock by GBM and LGG. Provided as a separate file.

